# 3D Analysis of the Force Generated by a Honeybee During Flight

**DOI:** 10.1101/2022.11.25.517967

**Authors:** Mandiyam Y. Mahadeeswara, Mandyam V. Srinivasan

## Abstract

To better understand the turning flight characteristics of the bees, we developed a procedure for estimating the instantaneous total force generated by the bee, along with its centrifugal force component, at each instant of time. We calculated the magnitude and direction of the total force vector (TFV) with respect to the three body axes of the bee and examined its variation with time during turning flights. The results of this study revealed that turns in the cloud are executed by (a) holding the magnitude of the TFV vector constant and (b) by redirecting the body (and therefore the turning force) appropriately to execute a coordinated turn. We also calculated and analysed the TFVs of bees flying in a curved tunnel. The characteristics of these TFVs (magnitude and direction) are very similar to those for bees flying in the cloud. This is a novel finding as there are no studies which have estimated the instantaneous flight force magnitude and direction of a turning bee (these are not saccadic or evasive turns) in an outdoor or indoor setup. However, similar results have been reported for pigeons and cockatiels while executing smooth turns in the horizontal plane.

## INTRODUCTION

Flying insects have the ability to perform a wide range of complex tasks –regulation of flight speed^1-3^, avoiding stationary^4-6^, moving^7,8^ obstacles, performing smooth landings^9,10^ and altitude^11^. Successful execution of these tasks requires subtle to significant changes in the direction and the magnitude of the instantaneous flight force generated by the insect. Understanding how an insect achieves this is of interest to biologists as well as engineers.

The magnitude and direction of the total force produced by a flapping insect is directly linked to the wing articulation and depends on a number of parameters, including the amplitude and frequency of the wing beat, the stroke plane, and the angle of attack of the wings in relation to the stroke plane. Several investigations have used high-speed filming to capture the wing kinematics and applied computational fluid dynamics to estimate the lift and thrust forces that are generated during flight. Classic studies from the laboratory of Goetz^12^, in which tethered, flying flies were exposed to stimuli moving in various directions with respect to the body (upward, downward, forward, backward, and obliquely) revealed that the direction of the flight force (which was measured by a force transducer coupled to the tethering device) was oriented in a fairly constant direction in relation to the longitudinal axis of the body, irrespective of the visual stimuli that were delivered. And this was true even when the tether was rotated to change the pitch of the body^13^. The orientation of this force was found to be inclined forwards and upwards by 24 deg with respect to the longitudinal body axis in *Drosophila*, and 29 deg in *Musca*^12^. While these are important findings, one cannot exclude the possibility of artefacts, induced by the tethering, that could alter the magnitude and direction of the measured flight force. It is therefore desirable to investigate the magnitude and direction of the total force that insects generate during free flight, and examine whether, and if so, how this force varies while the insect performs manoeuvres that include roll, yaw, and pitch.

In this study we address this question by filming honeybees during free flight in the outer periphery of the cloud and in the curved tunnel, and analysing the video images to measure the instantaneous speed and direction of flight, as well as the instantaneous pitch and roll angles during various turning manoeuvres and in straight flight. This data is used in combination with knowledge of the honeybee’s mass, estimates of the passive lift and drag forces generated by the body, and computations of the total instantaneous acceleration generated by the bee (the vector sum of the tangential and the centripetal accelerations), to estimate the magnitude of the total force generated during flight, and the direction of this force with respect to the longitudinal axis of the bee.

## METHODS

### RECORDING AND RECONSTRUCTING TURNING FLIGHTS IN THE PERIPHERY OF THE BEE CLOUD

Two downward-facing cameras were used to reconstruct the 3D flight trajectories of bees on the outer periphery of the bee cloud arena, as shown in Fig. 1a. The video frames from the two cameras were analysed to reconstruct the 3D flight trajectories, and to measure the instantaneous body orientations of bees flying in the cloud, as described in this study^14^. However, capturing the roll angles of a bee flying in a cloud is more challenging. This is because the flights are in different directions and in different planes, so that recording the roll angles of a bee over a continuous sequence of frames is a challenging and arduous task, in relation to camera positioning. However, for the purposes of the present study, we were able to measure the trajectories and roll angles for a few flights in the periphery of the bee cloud, using appropriate camera positioning. The cameras for measuring the roll angles were mounted horizontally (i.e. with their optical axes in the horizontal plane). As a consequence, the segments of the flights that were analysed were confined approximately to the horizontal plane.

**Figure 1:**
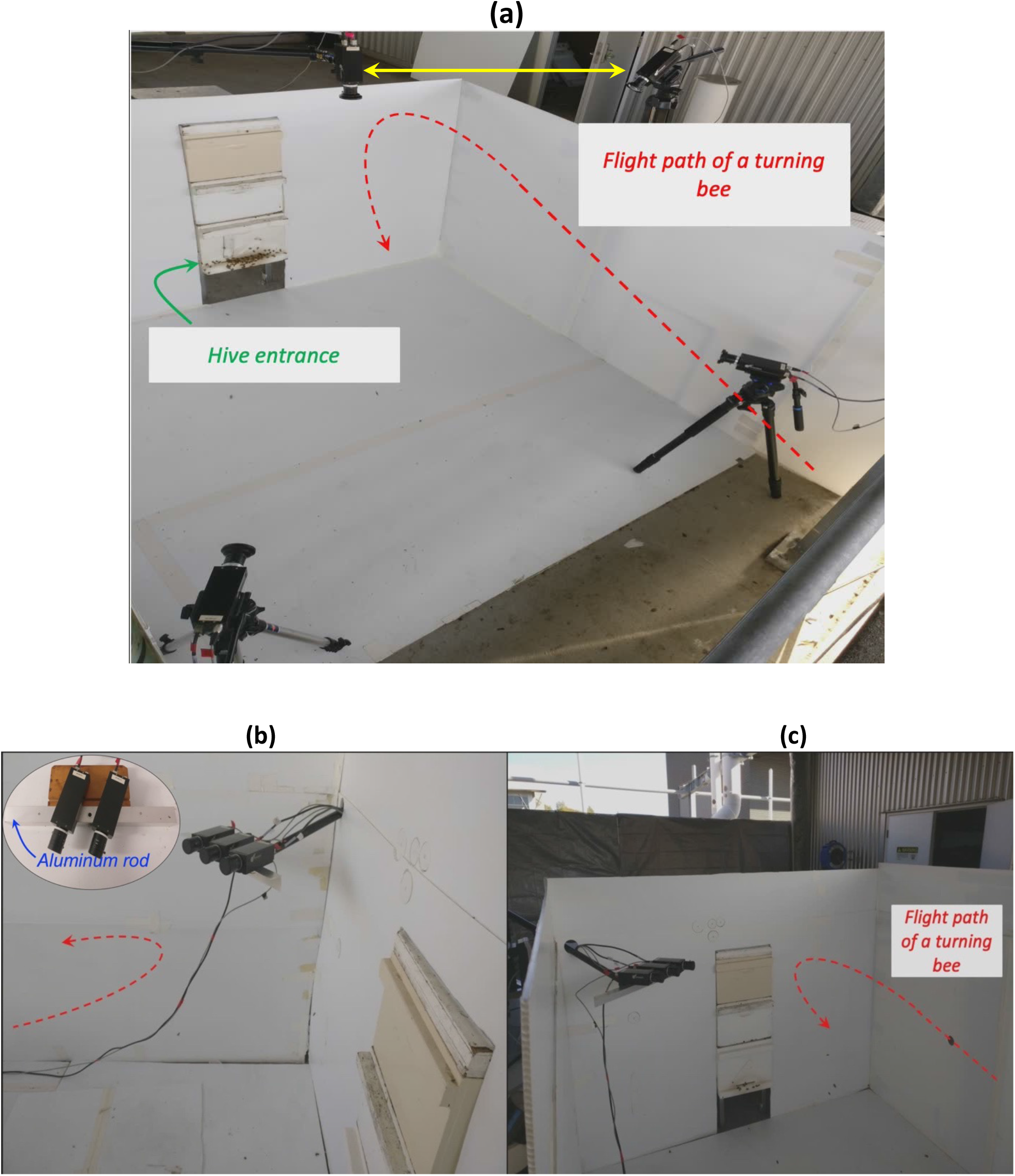
(a) An image of the experimental arena on our building rooftop showing the flight path of a turning bee (dashed red line). The double arrow in yellow shows the locations and orientations of the two cameras used to record the overhead view of a turning bee. (b,c) Images showing the camera arrangement employed to capture the frontal view of a turning bee. The reason for positioning two frontal-view cameras next to each other is to obtain a greater depth of field and capture a longer flight segment. One camera would cover the region where the bee is making a turn, and the other camera would cover the segment ahead of the turning region. The inset in panel (b) shows a close-up view of the camera configuration for capturing roll.

In order to capture a continuous sequence of roll angles made by a turning bee in this flight path, one has to (a) position the cameras in such a way that the flight of the bee is not disturbed; (b) adjust the positioning and optics of the cameras so that the bee is continually in the camera’s field of view and in focus in the turning region and to orient the cameras so that their optical axes are approximately parallel to the bee’s body axis in 3D. In other words, one should be able to see only the head and wings of the turning bee in the camera image. In this way, one can obtain reliable measurements of the roll angles.

A fully accurate measurement of the roll angle is obtained if the optical axis of the camera is exactly parallel to and collinear with the body axis. It is practically difficult (if not impossible) to meet this requirement for every video frame, because the cameras are fixed and the bees are continually in motion. However, small mismatches in the orientations are likely to lead to very small errors: for an angular discrepancy of δ between the body axis and the camera axis, the error in the measured roll angle is proportional to 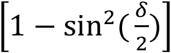.

To capture the roll angles, we mounted two cameras next to each other on a custom-built aluminum bar (inset, Fig. 1b) mounted on a tripod. The reason for using two cameras with parallel optical axes was to obtain a higher depth of field while recording the roll angles. The cameras were positioned in the experimental arena as shown in Figs. 1b, c. (three cameras are shown in Figs. 1b, c but in practice only two were used to record the roll angle information, as shown in the inset in panel 1b). Using arrangement, we were able to successfully record roll angle information for short durations of time while the bees were executing turns in the region covered by the cameras.

Using the above procedure, a total of 27 flight trajectories were recorded, along with roll angle information. It is important to note that the roll angles of a turning bee are not available for the entire turning sequence, because the bee’s heading changes gradually during the smooth turning flights. There would be a point where the optical axis of the camera recording the roll angle is not parallel to the bee’s body axis. In these circumstances, the digitised roll angles will not be an accurate measure of the roll angle held by the bee. Consequently, roll angles were not always available for the entire flight segment that was captured.

### RECORDING AND RECONSTRUCTING FLIGHT TRAJECTORIES IN THE CURVED TUNNEL

The experimental setup for the tunnel study is shown in Fig. 2(a-b). The tunnel (see Fig. 2(a)) consists of four sections: Section 1: Straight; Section 2: Curved; Section 3: Straight; Section 4: Slightly curved.

**Figure 2:**
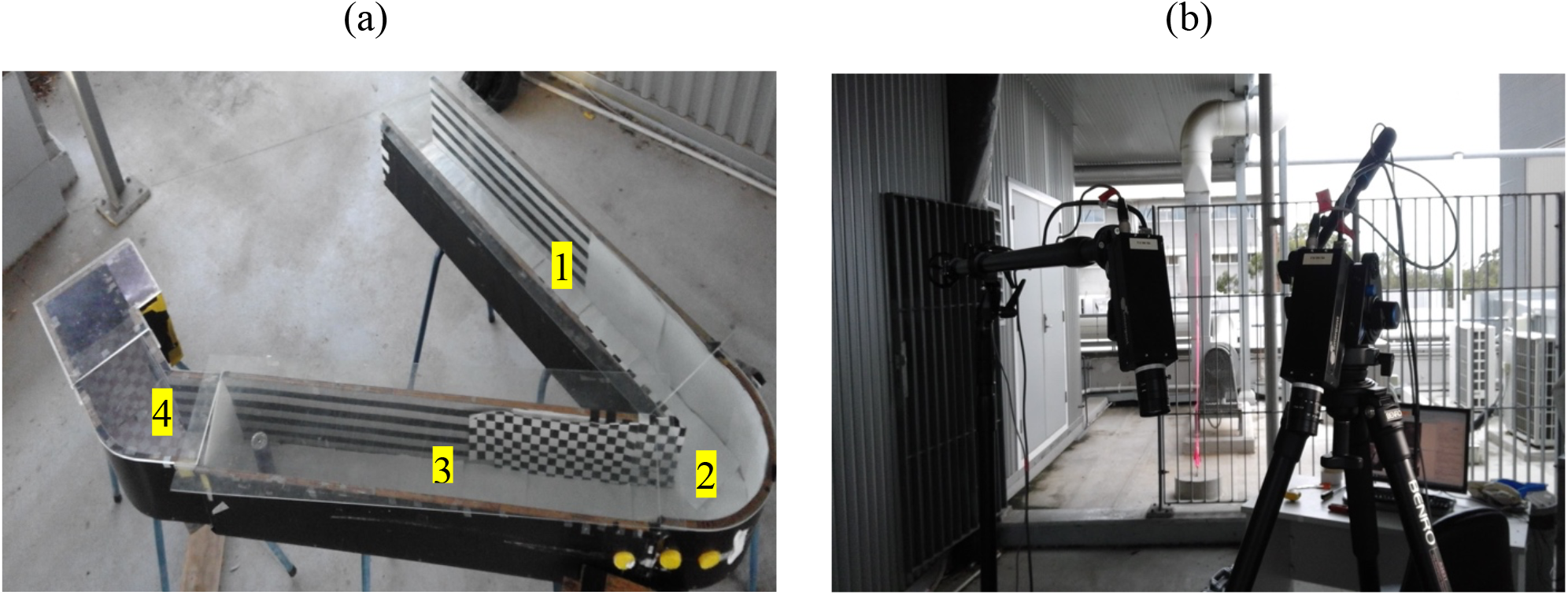
(a) View of tunnel comprising a curved section (2) connecting two straight sections (1,3). (b) View of overhead high speed video cameras for filming and reconstructing the bee trajectories in 3D.

The length, width, and height of the straight sections of the tunnel were 1400 mm, 230 mm and 194 mm, respectively. The width and height of the curved section of the tunnel were 230 mm and 194 mm, respectively. The radius of curvature of the mid-line of the curved section was 115 mm. Schematic 3D and plan views of the curved section of the tunnel is shown in Fig. 3.

**Figure 3:**
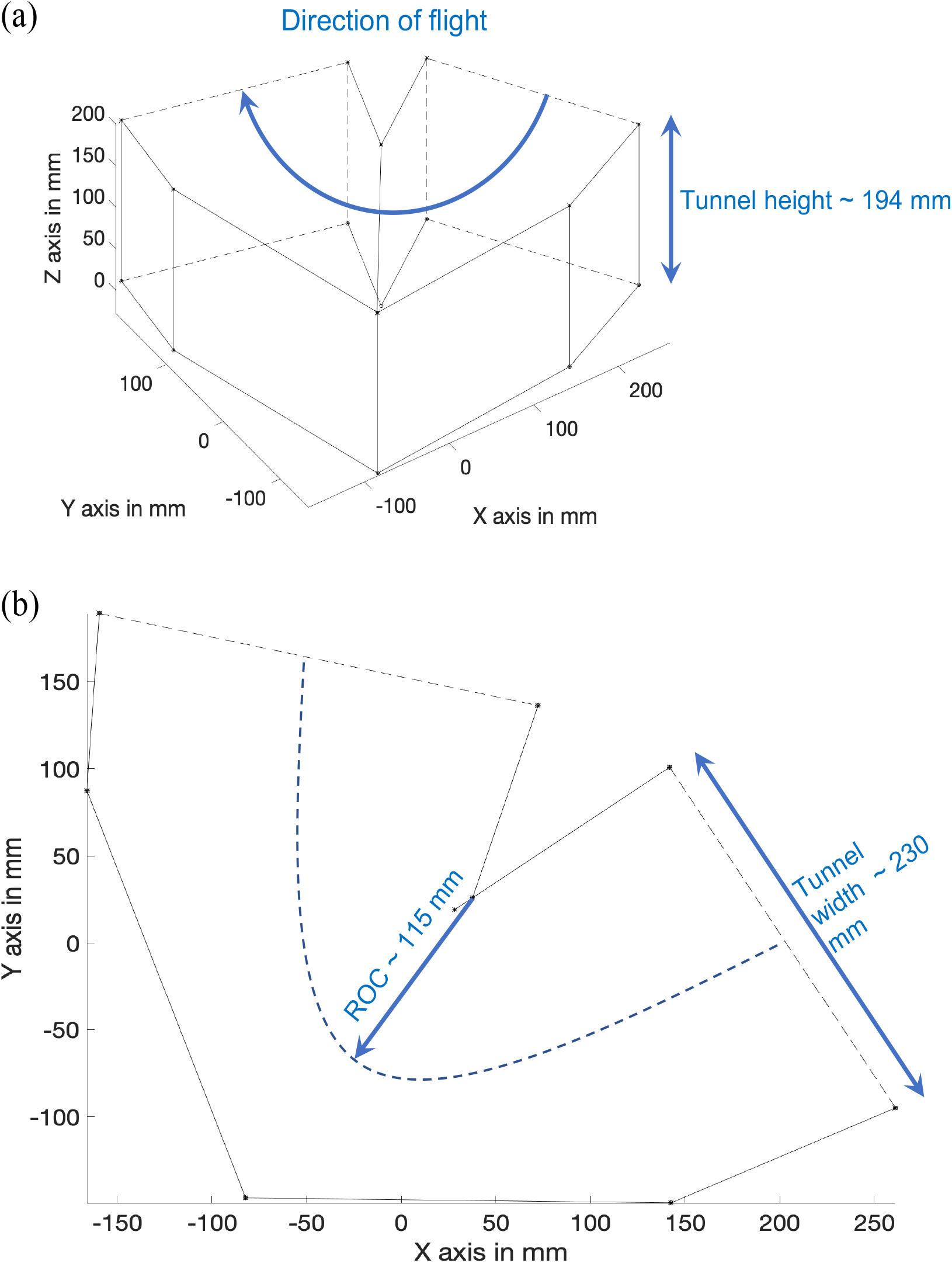
(a) A schematic 3D view of the curved section of the tunnel (b) A plan view of the curved section showing the relevant dimensions. (The outer wall is actually curved, as shown in the image of Fig. 4(a), with a radius of 230 mm, but it is shown here as being composed of two planar surfaces, for simplicity of illustration).

Only flights in the curved section (Section 2) were considered for this analysis, as the main aim was to investigate turning behaviour. The experiment was carried out in a semi-outdoor environment. We commenced the experiment by training the bees to fly into the tunnel by placing a sugar water feeder at the tunnel entrance and then slowly moving the feeder into and along the tunnel, all the way to its far end. It took about a day to train the bees to fly smoothly in and out of the tunnel. We allowed another half a day for the bees to get acquainted with the tunnel environment.

Flights in the tunnel were recorded by a high-speed camera system (Emergent Vision Technologies) consisting of 4 synchronized cameras with 2048 ×1080 pixel resolution, running at 335 fps.

Two cameras, positioned above the tunnel (Fig. 2b), were used to capture and reconstruct the 3D trajectories of bees flying through the tunnel. The tracking and stereo matching algorithms explained in the study^14^ were used to obtain the final 3D head and tail positions of bees flying through the tunnel. 37 bee flights in the tunnel were digitised and reconstructed for analysis. Only those trajectories in which a bee flew in isolation (without interference from other bees flying in the same or the opposite direction) were considered for analysis.

### MEASUREMENT OF ROLL ANGLES IN THE CURVED TUNNEL

Two additional cameras, viewing the tunnel through apertures in the side wall (Fig. 4a) were used to capture frontal (head-on) views of the bees, to measure their roll angles as they were making their turns (Fig. 4b).

**Figure 4:**
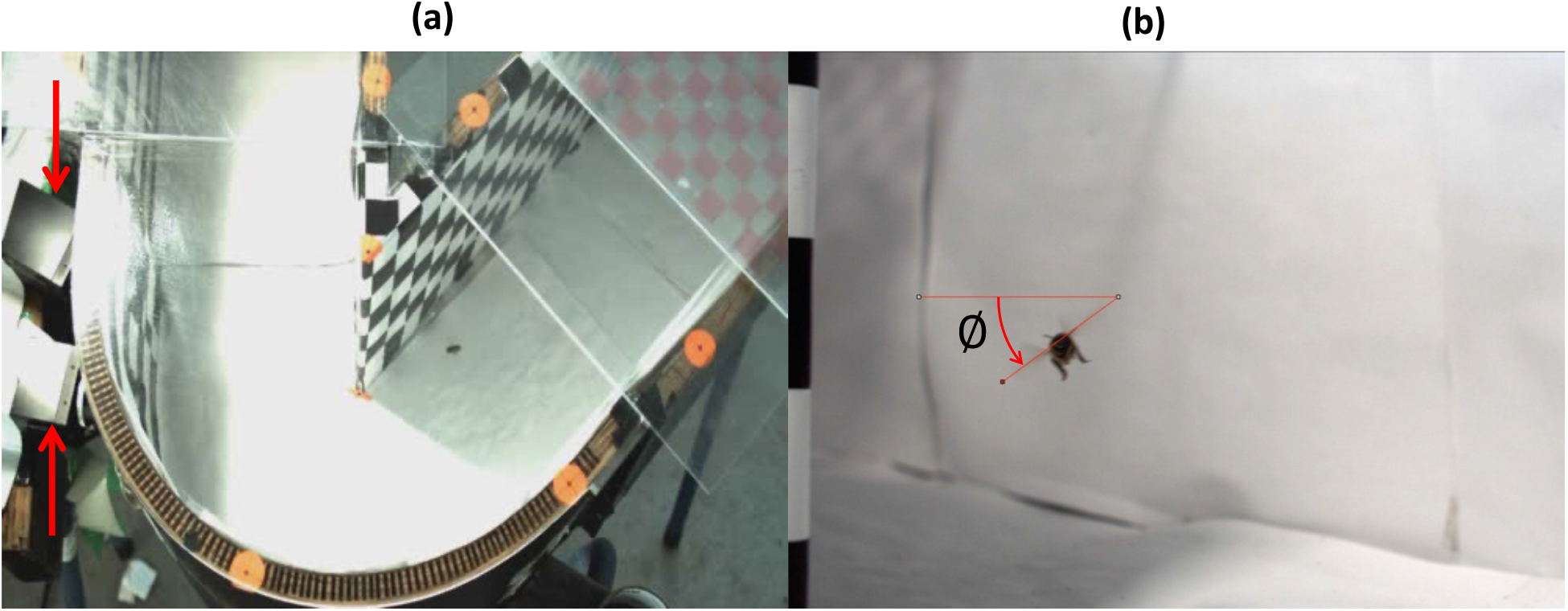
(a) Close-up view of a bee making a turn in the tunnel, along with the cameras (red arrows) used for recording the roll angles (b) Head-on view of a bee making a turn in the tunnel. The roll angle (∅) is measured using the angle tool in ImageJ.

The roll angle of the bee was measured from the frontal view videos by manually digitising the pixel locations of the two wing tips and using ImageJ to measure the orientation of the line connecting the two wingtips with respect to the horizontal axis, as shown in Fig. 4b. Making these measurements from the side views with good resolution and depth of field at high frame rates was a challenging task. It entailed a lot of preliminary testing with various visual backgrounds on the inside tunnel walls, in order to clearly visualize the insect’s wing tips.

### ESTIMATION OF THE TOTAL FLIGHT FORCE GENERATED BY THE BEE DURING FLIGHT IN THE HORIZONTAL PLANE

The procedure for estimating the total flight force is illustrated in Fig. 5, which shows (a) the force required to support the weight of the body (acting vertically upwards, blue arrow); (b) the force required to generate the tangential acceleration (pointing along the flight direction, dashed red arrow); (c) the force required to oppose the centrifugal force (the centripetal acceleration, pointing toward the centre of curvature, dashed red arrow); (d) the total force (solid amber arrow) required to oppose the body-induced drag (pointing opposite to the flight direction, dashed amber arrow) and the body-induced lift (pointing vertically upwards, dashed amber arrow). These vectors are represented with respect to the external world coordinates in Fig. 5. The sum of all these vectors (shown in green) is the total flight force vector, which specifies the magnitude and direction of the instantaneous total flight force, in external world coordinates.

**Figure 5:**
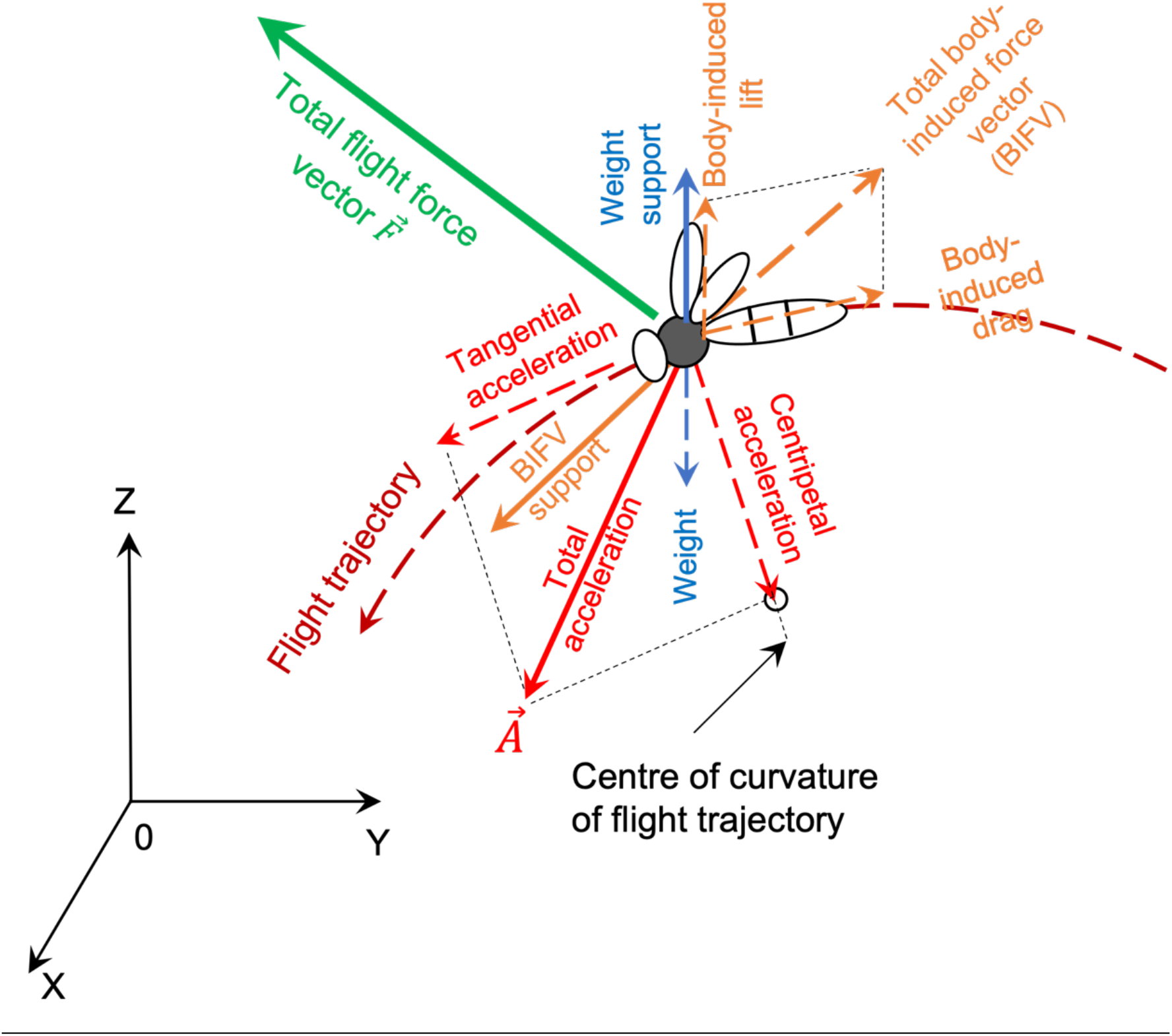
Diagram showing an oblique view of a bee flying along a curved trajectory, illustrating the various components that make up the total flight force.

### ESTIMATION OF THE BODY-INDUCED FORCE VECTOR

The magnitudes of the body-induced drag and lift forces generated by the movement of the bee in still air depend upon (i) the flight speed and (ii) the pitch angle of the body with respect to the flight direction, as demonstrated by Nachtigall and Hanauer-Thieser^15^. The body pitch angle decreases with increasing flight speed. The authors measured the lift and drag forces experienced by a tethered bee when it was exposed to four different, calibrated air speeds in a wind tunnel, with the body oriented at four different pitch angles relative to the direction of the air flow. An example of their results is illustrated in Fig. 6 (adapted from Fig. 6B of their paper), which shows the drag (D) and lift (L) forces, and the magnitude and the direction of the resultant, total body-induced force vector (which is termed as BIFV), for an airspeed of 1 m/s and a body pitch angle of 20 deg. In this case the drag force is 1.73 ×10^−5^ N, and the lift force is 3.60 ×10^−6^ N. The total, resultant body-induced force has a magnitude of 1.77 ×10^−5^ N and is inclined at 11.8 deg relative to the (negative) flight direction.

**Figure 6:**
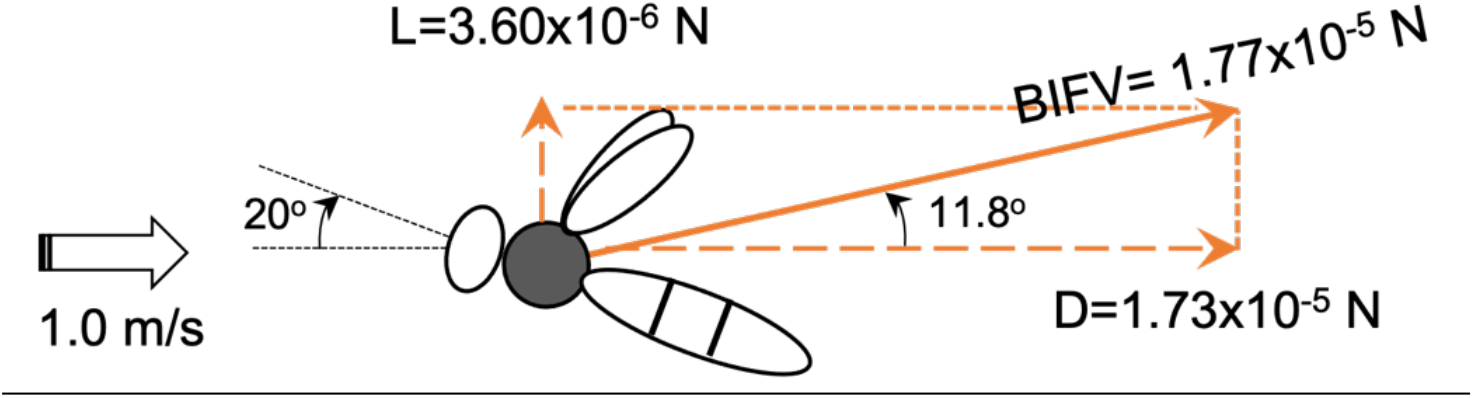
An example of the body-induced drag (D), lift (L) and total force (BIFV) at a pitch angle of 20° and a horizontal airspeed of 1m/s, as reported by Nachtigall and Hanauer-Thieser^15^.

The complete set of results obtained by Nachtigall and Hanauer-Thieser^15^ is summarised in Table 1. We used these published measurements of the BIFV to estimate the magnitude and direction of the total flight force generated by the bee at each instant during its flight, based on my measurements of the flight speed and body pitch from the videography of the bees’ flights.

**Table 1:** Magnitudes and directions of the body induced force vector (BIFV) experienced by a bee at different airspeeds and pitch angles (reproduced from Nachtigall and Hanauer-Thieser^15^)

|  |  |  |  |  |
| --- | --- | --- | --- | --- |
| **Airspeed (m/s)** | 0.25 | 1.0 | 5.0 | 12.0 |
| **Body pitch angle (deg)** | 45 | 20 | 5 | 0 |
| **BIFV orientation** [Angle relative to flight direction (deg)] | 17.9 | 11.8 | 14.6 | 7.3 |
| **BIFV magnitude (N)** | $1.61 \times 10^{-6}$ | $1.77 \times 10^{-5}$ | $1.72 \times 10^{-4}$ | $1.15 \times 10^{-3}$ |

We used interpolation to estimate of the magnitude and direction of the BIFV for airspeeds and body pitch angles not shown in Table 1. The orientation of the BIFV for a measured body pitch angle that was not in Table 1 (but was within the range spanned by the data therein) was estimated through linear interpolation of the BIFV orientation values for flanking body pitch angles in the table. Thus, for example, the BIFV orientation [denoted by ∠*BIFV*(10 *deg*)] for a measured body pitch angle of 10 deg was calculated by linearly interpolating the BIFV orientations given in the table for the flanking body pitch orientations of 5 deg and 20 deg, as

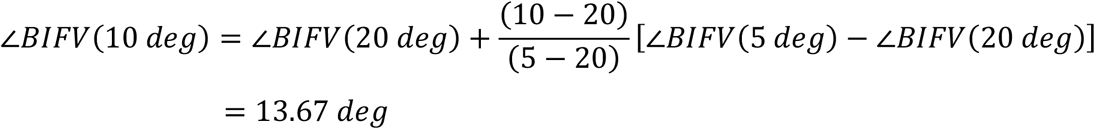

This linear interpolation yields fairly accurate estimates, as is evidenced by the data shown in Fig. 6A of Nachtigall and Hanauer-Thieser^15^. However, linear interpolation does not provide accurate estimates of the *magnitude* of the BIFV across various airspeeds, because drag forces typically increase approximately as the *square* of the airspeed^15^. Therefore, estimates of the magnitude of the BIFV for a measured flight speed that was not in the table (but was within the range spanned by the data therein) were obtained using quadratically extrapolated values from the BIFV magnitudes given in the table for the two flanking airspeeds, and averaging the two extrapolated values. Thus, for example, the BIFV magnitude [denoted by |*BIVF*|] for a measured flight speed of 3m/s was estimated from the tabled values for the BIFV magnitudes at 1.0 m/s and 5.0 m/s as:

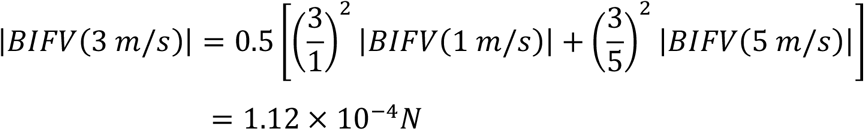

#### ESTIMATION OF THE FORCES REQUIRED TO GENERATE THE ACCELERATIONS

In brief, the force vector describing the instantaneous total acceleration 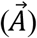 due to the bee’s motion is computed for each video frame as 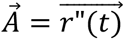 where 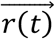 is the 3D vector representing the instantaneous position of the bee (in an external coordinate frame), as described in the study by Mahadeeswara and Srinivasan^16^. The tangential and centripetal components of this force vector (if required, for additional information) is computed as

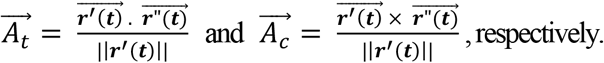

### ESTIMATION OF THE TOTAL FLIGHT FORCE VECTOR

The total flight force vector is estimated by summing its various components, which are illustrated in Fig. 5. The total flight force vector 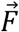 (expressed in the external coordinate frame) is the vector sum of the forces required to generate the total acceleration, to compensate for the body weight, and to compensate for the body-induced drag and lift forces (noting that the body-induced lift partially compensates for the body weight):

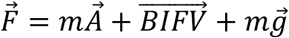

where *m* is the mass of the bee and 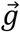 is the (negative) acceleration due to gravity, representing the upward acceleration required to compensate the bee’s weight. The bee’s body weight is assumed to be 0.1 grams.

### PROCEDURE FOR CONSTRUCTING THE REFERENCE BODY AXES OF THE BEE

The above calculations deliver the magnitude and direction of the total flight force F in the external coordinate frame, and not relative to the bee’s body orientation. In order to determine the orientation of the total force vector 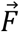 with respect to the three body axes of the bee, one has to construct the three body axes by including the attitude of the bee in the pitch, roll and yaw axes. Using the bee’s head-to-tail vector (obtained by computing the 3D coordinates of the head and the tail from the digitised video frames), we first define an axis that points in the direction of the bee’s longitudinal body axis. We call this axis the ‘frontal axis’ of the bee, and denote it by the unit vector 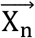 where n denotes the frame number).

The unit vector 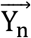, defining the lateral axis of the bee’s body (pointing toward the bee’s left) is then given by:

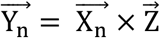

where 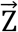 is the unit vector directed along the vertical axis of the external co-ordinate system, and × denotes the cross product.

Finally, we compute the dorsal axis of the bee (denoted by the vector 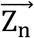), between X_n_ and Y_n_:, by performing a cross product

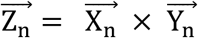

We have now constructed the three body axes of the bee, which take into account the rotations due to pitch and yaw. But the constructed axes should also include the rotation due to roll. To account for roll, the lateral axis 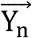 and the dorsal axis 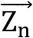 have to be rotated about the frontal axis by an amount equal to the roll angle, defined here as *θ*, to obtain the final orientations of the three body axes.

Techniques for rotating 3D coordinate axes are well established^17^. The procedure for rotating a 3D coordinate system by *θ* deg about an arbitrary, specified axis is as follows:

If the unit vector pointing along the axis of rotation is

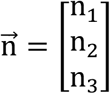

the rotation matrix R_n_(θ) is given by

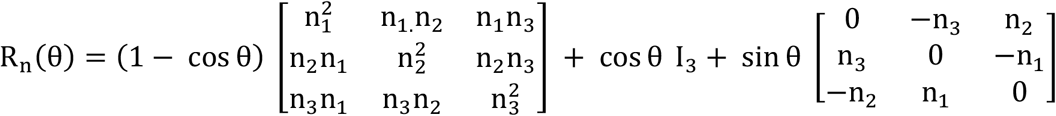

where I_3_ is a 3 × 3 identity matrix.

This matrix can be applied to rotate any arbitrary vector, say 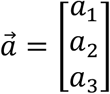 by premultiplying 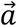 by *R*_*n*_(*θ*):

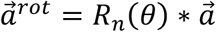

where (*) denotes matrix multiplication.

In our case, the rotation axis 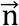 is the unit vector representing the frontal body axis of the bee 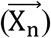, and the angle of rotation is *θ*, the roll angle. An illustration of how the three reference axes are reoriented after performing the rotation about X_n_ is shown in Fig. 7.

**Figure 7:**
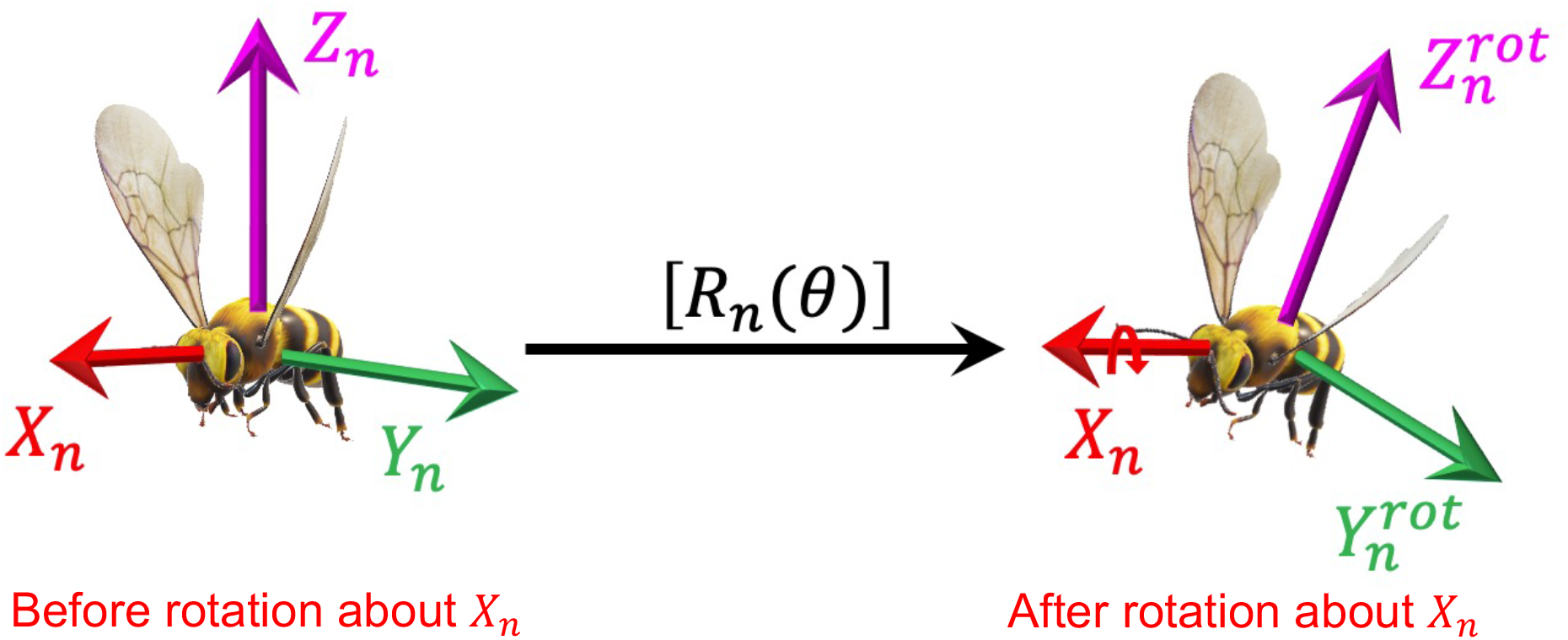
An oblique view of a bee showing the three reference axes before and after a rotation about the X_n_ axis (a counterclockwise roll). Clockwise roll angles are considered positive, in accordance with the right-hand system of coordinates.

### REPRESENTING THE TOTAL FLIGHT FORCE VECTOR IN THE REFERENCE BODY AXES OF THE BEE

The final step of the calculation is to determine the orientation of the total flight force vector 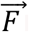 in relation to the bee’s body frame. Fig. 8 illustrates the axes of the external coordinate system (X, Y, Z) and the rotated reference body axes (x,y,z) of the bee, which have been computed as described above. We denote the unit vectors along x, y and z by 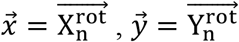 and 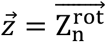, respectively. The orientation of 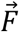 in the bee’s body frame is determined by computing the angles between 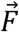 and the three body axes x, y, and z as 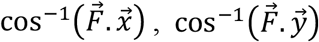 and 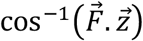, respectively.

**Figure 8:**
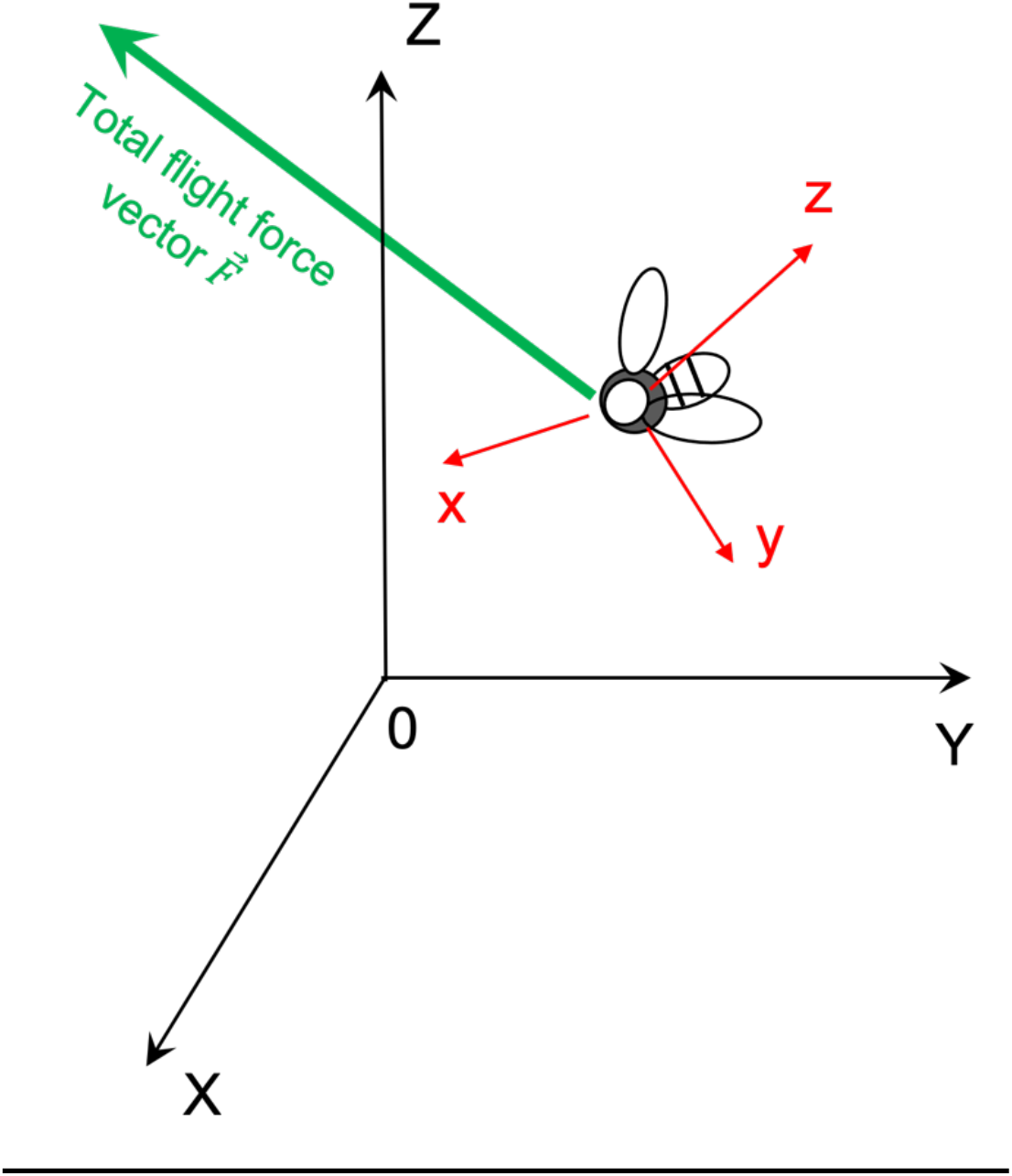
Representation of the total flight force vector in the reference axes of the bee’s body.

## RESULTS

### ESTIMATION OF THE TOTAL FORCE VECTOR FOR FLIGHTS IN THE CLOUD

We recorded flight segments of 27 different bees in the cloud, and used the procedure described above to estimate the total force vector (TFV) produced during each of these segments. Fig. 9a shows the magnitude of the estimated TFV (yellow curve) for each frame of a flight segment filmed for bee # 9. In this particular example, the magnitude of the TFV is (10.1± 0.9)×10-4 N (mean ± S.D). The variation (S.D.) of the TFV magnitude is less than 10% of its mean value, indicating that the total flight force is more or less constant throughout the flight segment. This is confirmed in the histogram of Fig. 10, which shows a tight distribution of the estimated TFV magnitudes around the mean value. The mean TFV magnitude (10.1×10-4 N) is only slightly greater (12%) than the weight of the bee (0.1 grams), which generates a downward force of (9.81×10-4 N). This is because a good part of the total force generated by the bee contributes to supporting its weight (which is compensated slightly by the upward body-induced lift).

**Figure 9:**
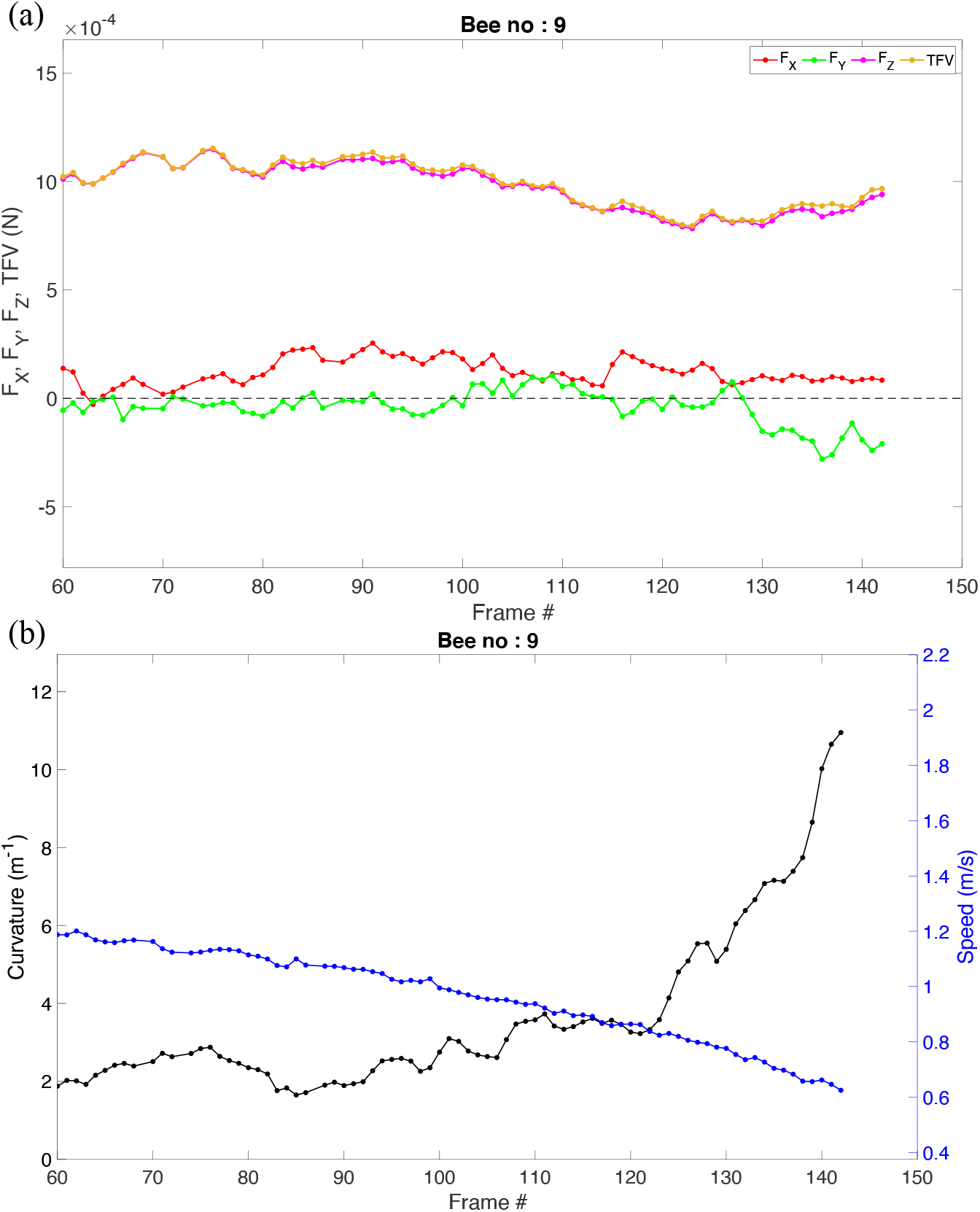
(a) The total force generated by bee # 9 during the analysed flight segment, along with its three components projected on to the reference axes of the bee. The total force is shown in yellow. The projections of the TFV on the frontal, lateral and dorsal axes of the bee are shown in red, green and magenta. (b) The blue and black curves show the variation of speed and curvature during the analysed flight segment.

**Figure 10:**
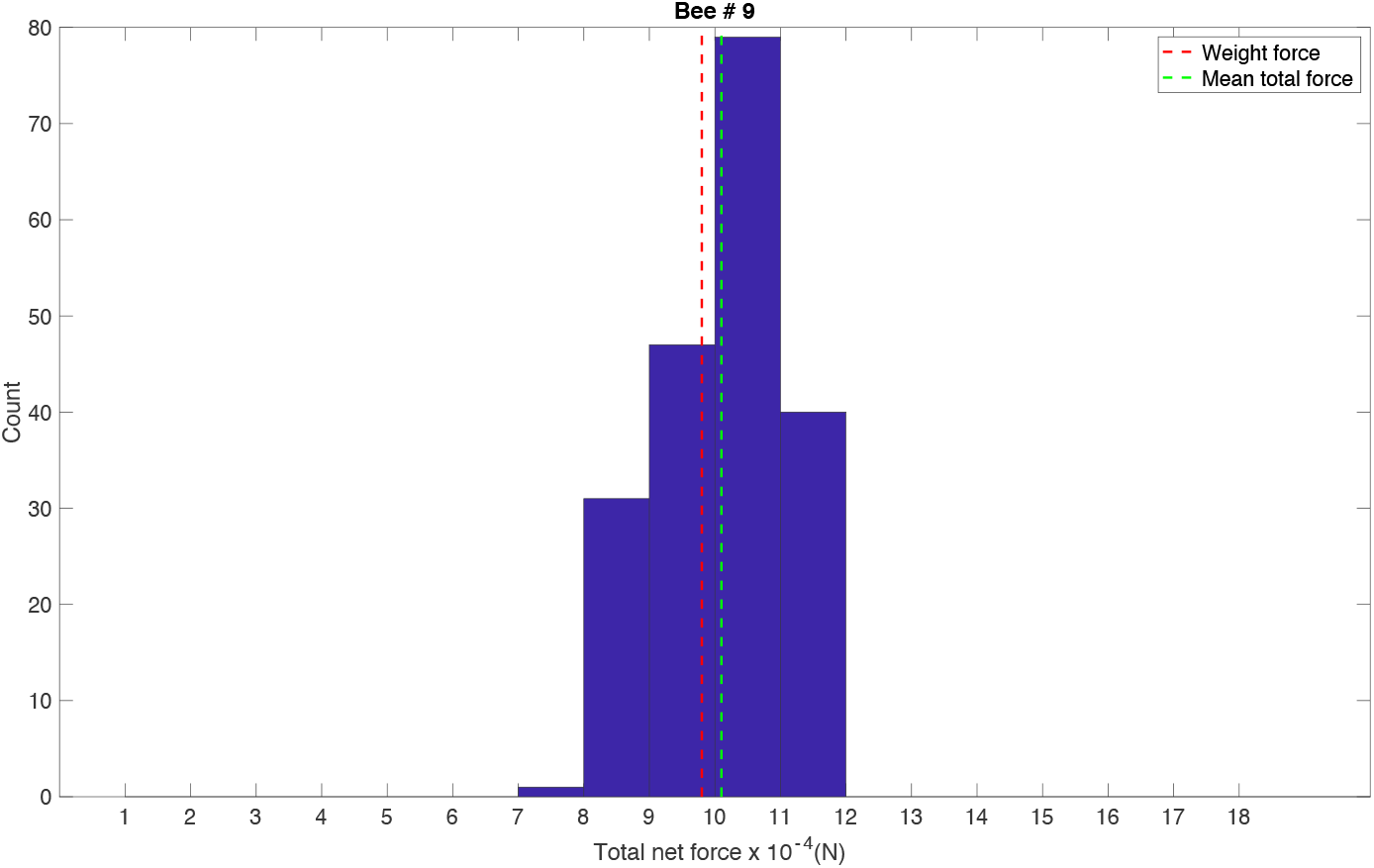
Histogram showing the distribution of the estimated magnitudes of the total force vector generated by bee #9 during the analysed flight segment. The dashed green line represents the mean magnitude of the TFV, and the dashed red line represents the downward force produced by the bee’s weight (9.81×10^−4^ N, assuming a bee mass of 0.1 grams)

**Figure 11:**
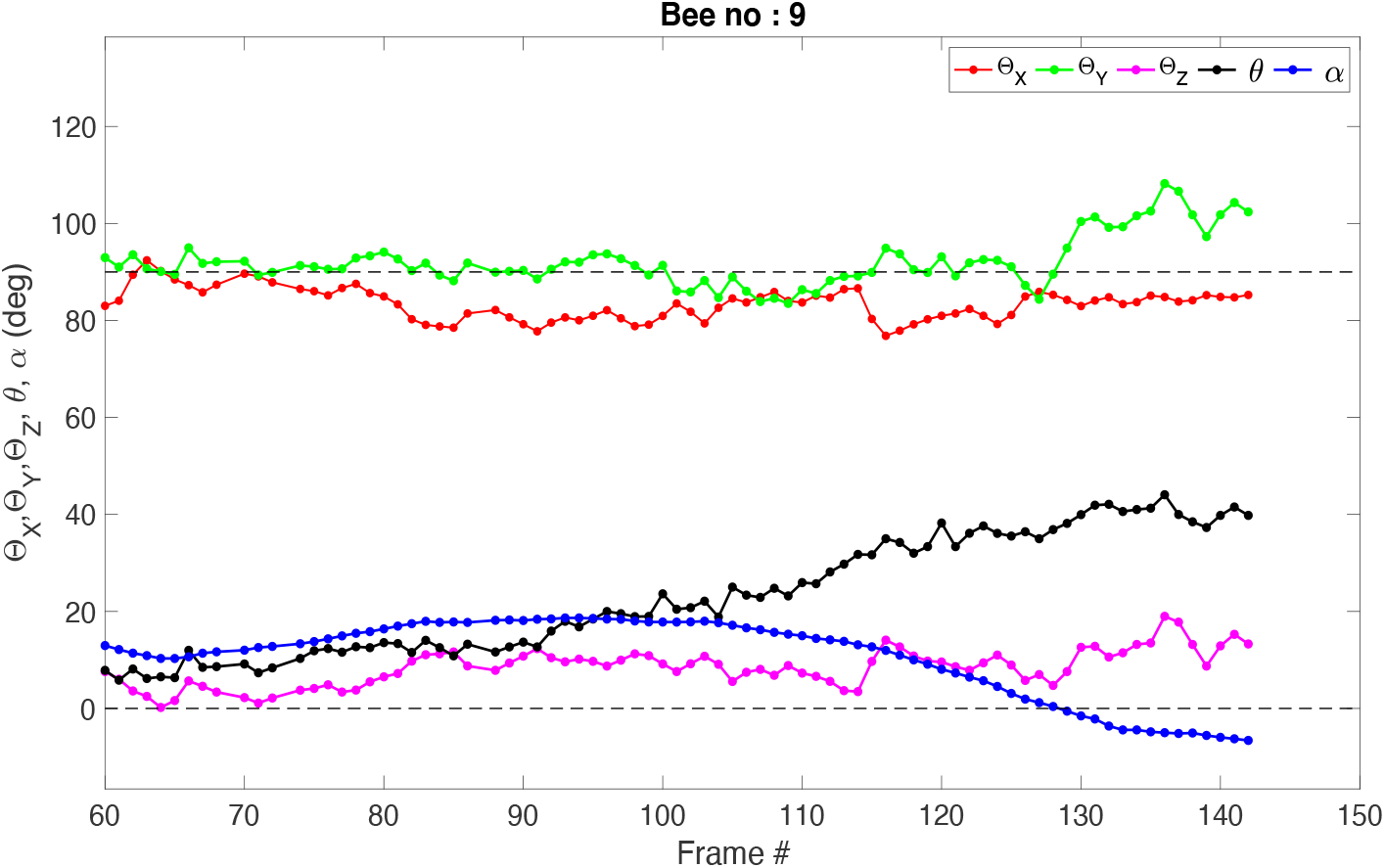
Variation of Θ_X_, Θ_Y_, Θ_Z_, θ(roll angle) and α(pitch angle) as a function of time for bee # 9 during the analysed flight segment. The profiles shown in red, green, magenta, black and blue represent Θ_X_, Θ_Y_, Θ_Z_, θ and α. Positive θ denotes a rightward roll.

Fig. 9a also shows the projections of the total force vector along each body axis of the bee. The major projection of the TFV is along the dorso-ventral axis (z axis) of the bee’s body (Fz). The small projection along the lateral (y) axis (F_Y_) indicates that the TFV is always nearly perpendicular to the bee’s lateral axis, and is not titled to the right or the left. This is true regardless of variations in the instantaneous speed or curvature of the flight trajectory, which are shown in the plots in the lower half of Fig. 9b.

Fig. 15 shows the orientation of the instantaneous total force vector with respect to the reference body axes of the bee in each frame of the flight segment. This plot reveals that the angles made by the estimated TFV vector with respect to the three body axes are always more or less constant, irrespective of changes in speed, curvature, or body orientation of the bee during the flight segment. For example, the angle (*Θ*_*X*_) between the TFV and the frontal body axis of the bee is held at a more or less constant value of 82.9 ± 3.2 deg. For the same bee, the angles made by the TFV with the lateral (*Θ*_*Y*_) and dorsal axis (*Θ*_*Z*_) are again more or less constant, at around 92.5 ± 5.1 deg and 8.9 ± 3.8 deg, respectively. The finding that *Θ*_*Y*_ exhibits a tight distribution around 90 deg indicates that the TFV is always nearly perpendicular to the bee’s lateral axis, and is not tilted to the right or the left.

The above observations are confirmed in Fig. 12, which shows histograms of *Θ*_*X*_, *Θ*_*Y*_ and *Θ*_*Z*_ all of which display tight distributions around their mean values.

**Figure 12:**
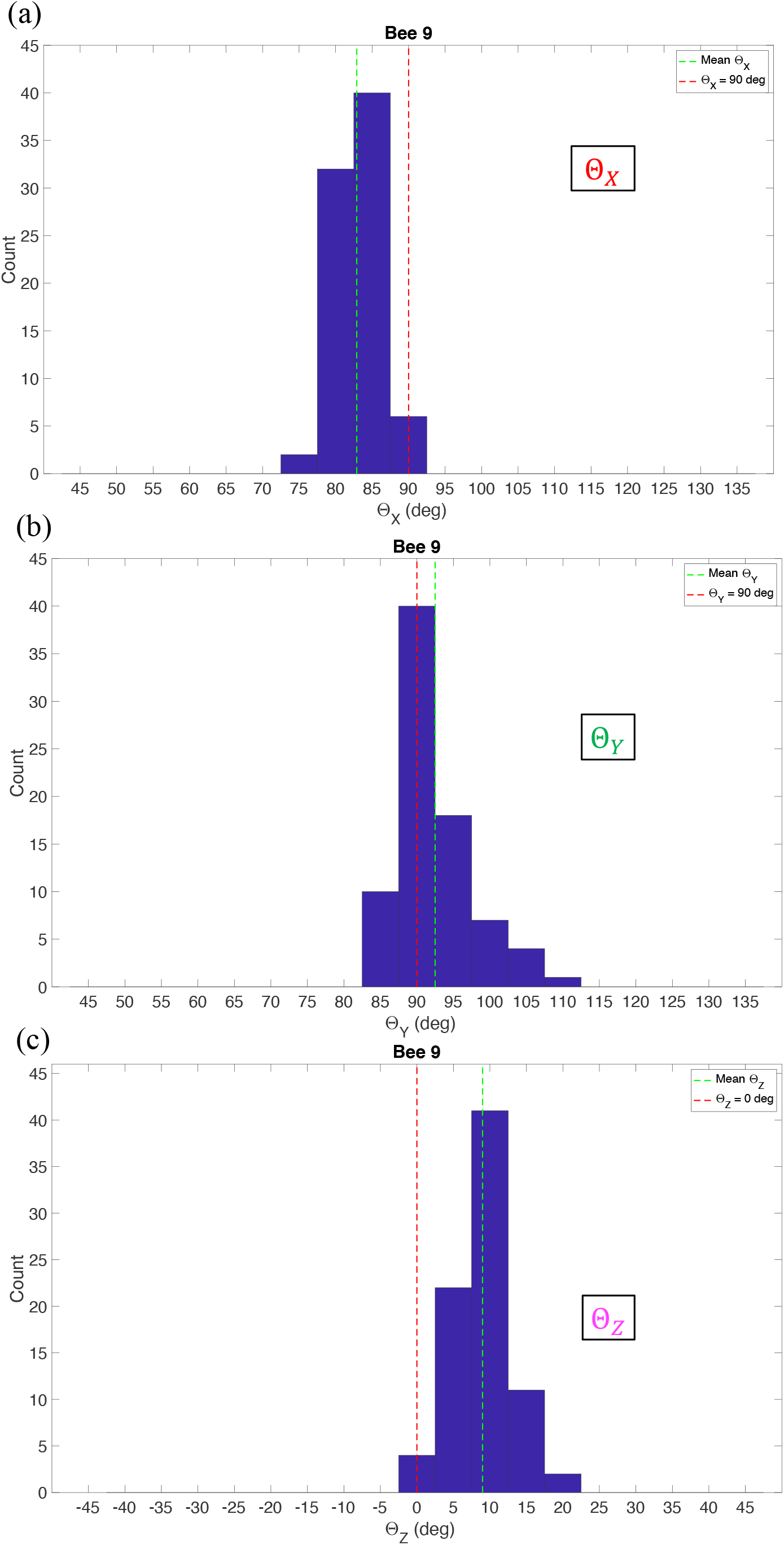
Histograms of Θ_X_, Θ_Y_ and Θ_Z_ for the flight segment of bee #9

Histograms of *Θ*_*X*_, *Θ*_*Y*_, and *Θ*_*Z*_, and the magnitudes and directions of the TFV for each of the 27 flight segments in the cloud that were filmed and analysed are given in Supplementary material (see Figs. S1). Table 2 shows the grand means of the various parameters that were measured and computed for these 27 flight segments.

**Table 2:** Grand means and standard deviations of Θ_X_, Θ_Y_, Θ_Z_, pitch angle (α), roll angle (θ), absolute roll angle, curvature magnitude, flight speed, and the magnitude of the total force vector (TFV) for the 27 flight segments filmed in the cloud. Most of the flight segments analysed in the cloud contained turns of the same polarity (right-hand turns, involving positive roll angles). Hence, the values of the mean and the SD for the (signed) roll angle are very similar to those for the absolute roll angle.

| $\theta_x$<br>(deg) | $\theta_y$<br>(deg) | $\theta_z$<br>(deg) | $\alpha$<br>(deg) | Roll<br>angle<br>(deg) | Absolute<br>roll<br>angle<br>(deg) | Tangential<br>acceleration<br>(m/s<sup>2</sup>) | Curvature<br>magnitude<br>(m<sup>-1</sup>) | Flight<br>speed<br>(m/s) | TFV<br>magnitude<br>(N) |
| --- | --- | --- | --- | --- | --- | --- | --- | --- | --- |
| 76.82<br>$\pm 8.33$ | 91.99<br>$\pm 3.02$ | 16.41<br>$\pm 6.91$ | 15.41<br>$\pm 4.70$ | 30.60 ±<br>9.51 | 30.75 ±<br>9.20 | -0.29 ± 1.86 | 8.09<br>± 3.88 | 0.93<br>± 0.17 | (11.1 ± 1.08)<br>$\times 10^{-4}$ |

The data in Table 2 confirm that (a) the magnitude of the TFV is maintained at a nearly constant value of (11.1 ±1.08)×10^−4^ N; and (b) the TFV vector is maintained at a more or less constant orientation relative to the body, irrespective of substantial changes in the speed, curvature and attitude of flight. Furthermore, the TFV is always oriented almost exactly perpendicular to the bee’s lateral axis; it is not tilted to the left or the right, irrespective of the attitude or speed of flight, or the curvature of the flight trajectory.

### ESTIMATION OF THE TOTAL FORCE VECTOR FOR FLIGHTS IN THE TUNNEL

We recorded flight segments of 37 different bees in the tunnel, and used the same procedure to estimate the total force vector (TFV) produced during each of these segments. Fig. 13a shows the magnitude of the estimated TFV (yellow curve) for each frame of a flight segment in the tunnel, filmed for bee # 5. In this case, the mean magnitude of the TFV is (9.68 ± 0.85)×10^−4^ N. The variation (S.D.) of the TFV magnitude is less than 10% of its mean value, indicating, again, that the total flight force is more or less constant throughout the flight segment, despite the variation in the instantaneous curvature (Fig. 13b). This is confirmed in the histogram of Fig. 14, which shows a tight distribution of the estimated TFV magnitudes around the mean value.

**Figure 13:**
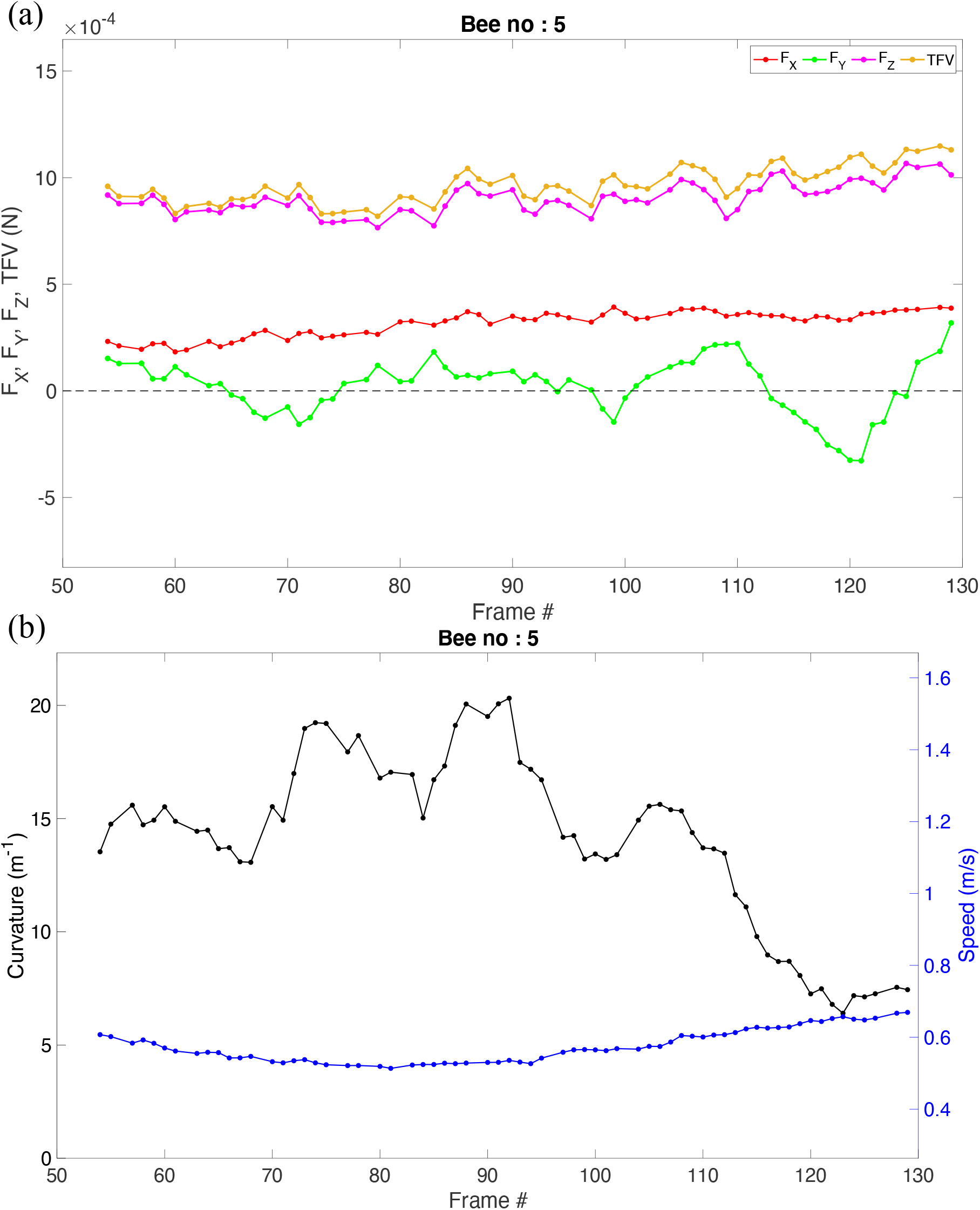
(a) The total force generated during a flight segment of bee # 5, along with its three components (F_X_, F_Y_ and F_Z_) projected on to each reference axis of the bee. The total force is shown in yellow. The projections of the TFV on the frontal, lateral and dorsal axes of the bee are shown in red, green and magenta. (b) The blue and black curves show the variation of speed and curvature during the analysed flight segment.

**Figure 14:**
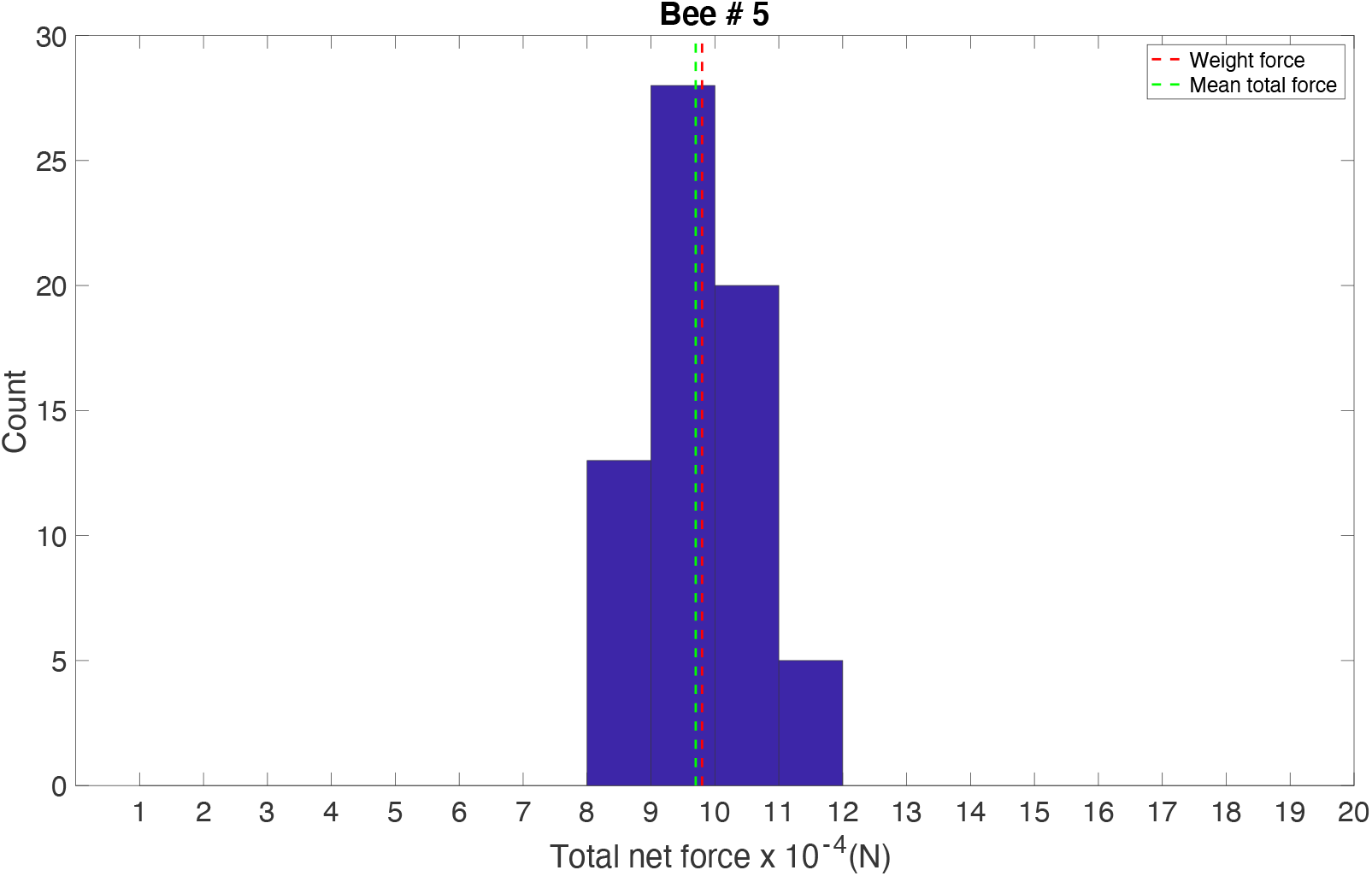
Histogram showing the distribution of the estimated magnitudes of the total force vector generated by bee #5 during the analysed flight segment. The dashed green line represents the mean magnitude of the TFV, and the dashed red line represents the downward force produced by the bee’s weight (9.81×10^−4^ N, assuming a bee mass of 0.1 grams).

**Figure 15:**
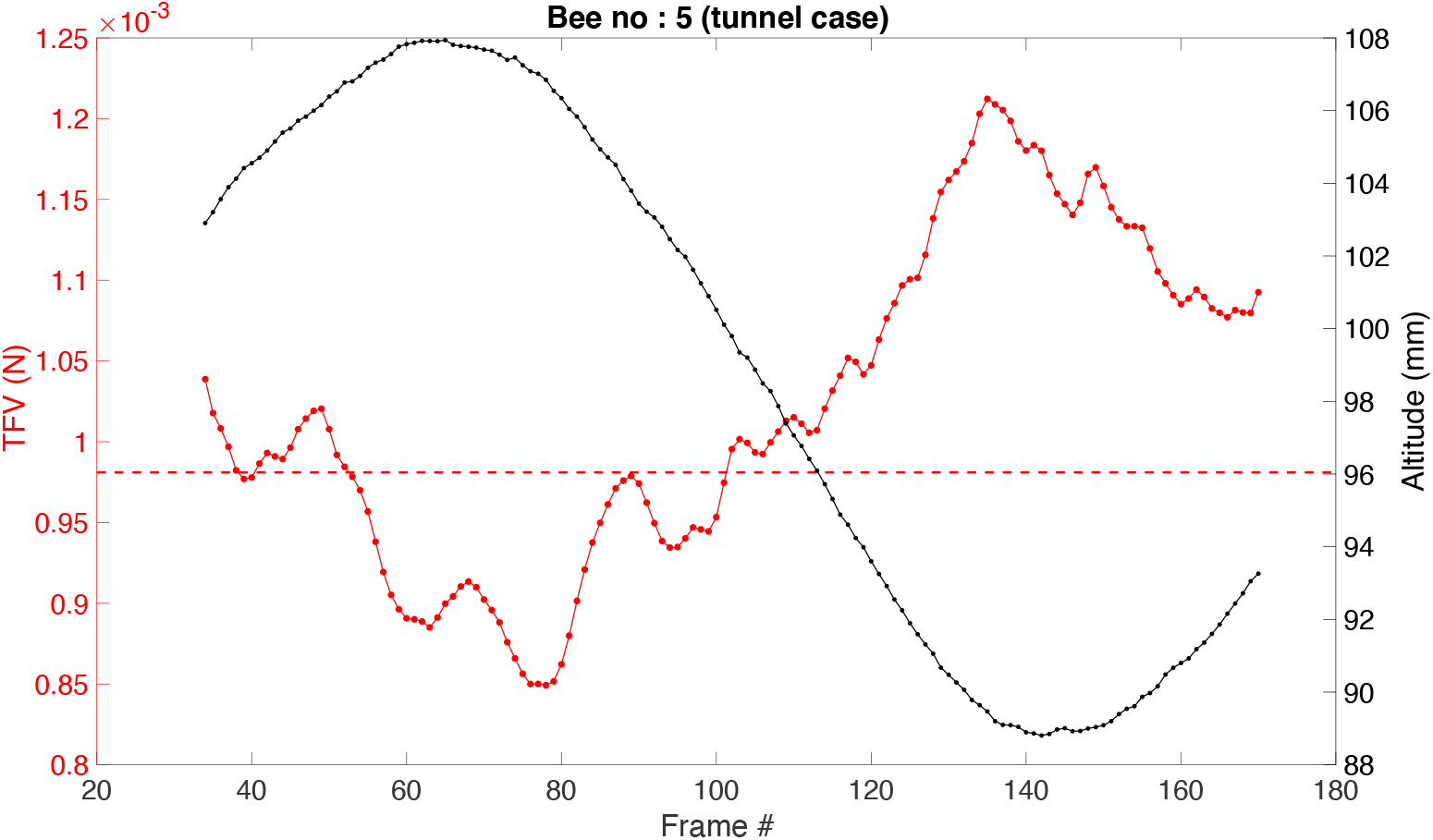
Variation of TFV magnitude and flight altitude for Bee #5. The flight segment spanning frames 54-129 is analysed in Figs. 13 and 14.

The mean TFV magnitude (9.68×10^−4^ N) is slightly lower than the downward force due to the weight of the bee (9.81×10^−4^ N). The reason for this may have to do with the accelerations that the bee undergoes in the vertical plane. This is illustrated in Fig. 15, which shows the variation of the TFV magnitude and the flight altitude during the flight segment. The height profile resembles that of a roller-coaster. The TFV magnitude is lowest (and below 9.81×10^−4^ N) in the vicinity of the crest of the height profile (frames 55-80), when the resulting upward centrifugal force partially cancels the downward force of the body weight. It is highest (and above 9.81×10^−4^ N) in the valley of the height profile (frames 130-160), when the resulting downward centrifugal force adds to the body weight. The variations in altitude are small, and the entire flight trajectory is mostly close to the horizontal plane – as is the case with most of the flight segments that were analysed.

Fig. 16 shows the orientation of the TFV with respect to the three body axes for the flight segment analysed in Figs. 13 and 14. The mean values and standard deviations of *Θ*_*X*_, *Θ*_*Y*_, and *Θ*_*Z*_ were found to be 70.9 ± 2.8 deg, 88.9 ± 7.6 deg and 20.6 ± 3.4 deg, respectively. The low standard deviations in all of these angles indicate that the orientation of the TFV is more or less constant in relation to the body, throughout the flight segment. These observations are confirmed in Fig. 17, which shows histograms of *Θ*_*X*_, *Θ*_*Y*_ and *Θ*_*Z*_, all of which display tight distributions around their mean values. The mean value of *Θ*_*Y*_ is very close to 90 deg, indicating that the TFV is oriented perpendicular to the lateral (y) axis, irrespective of the instantaneous attitude or speed of flight, or curvature of the flight trajectory. These findings are very similar to those obtained for the flight segment in the cloud (Figs. 9-13).

**Figure 16:**
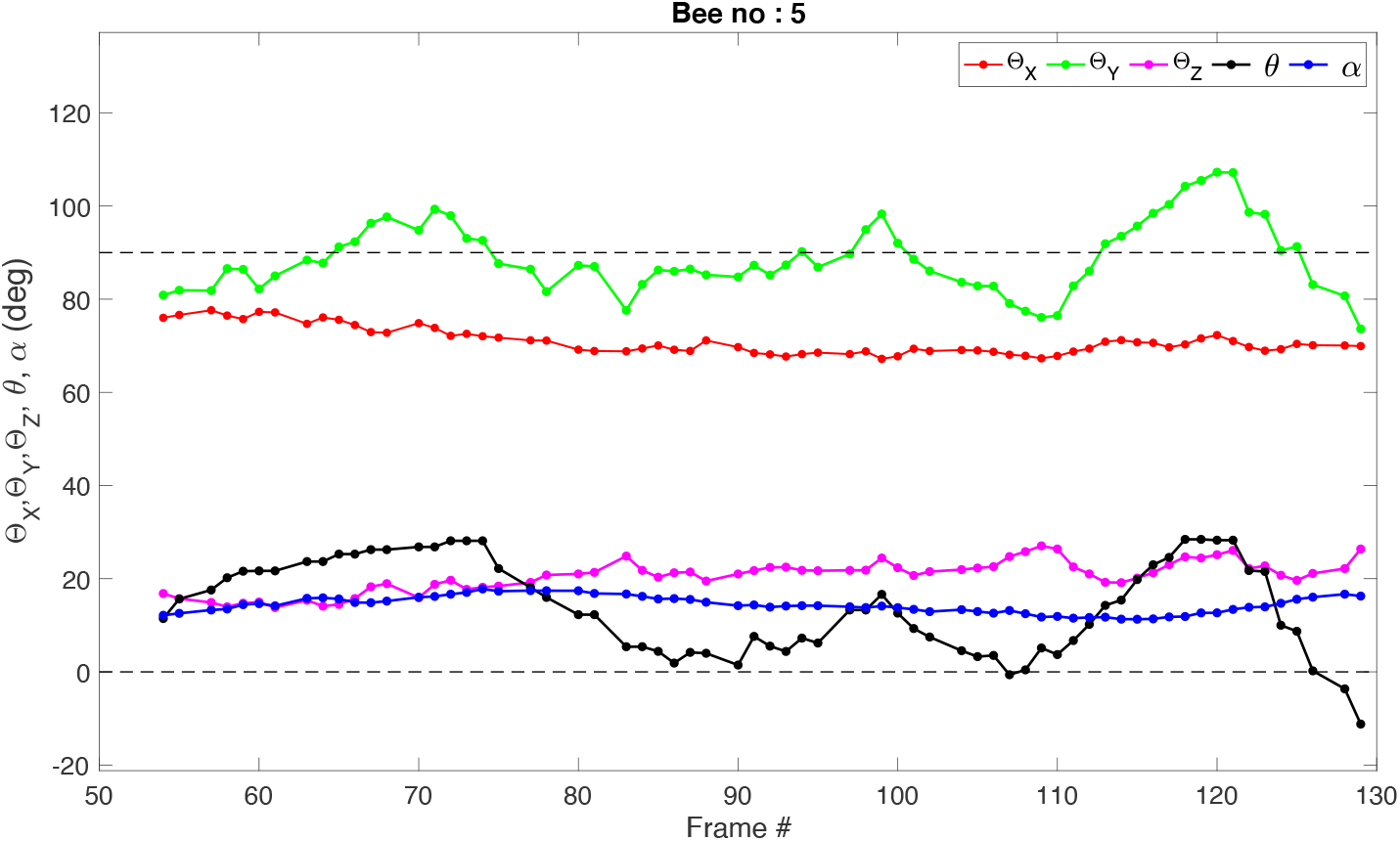
Variation of Θ_X_, Θ_Y_, Θ_Z_, θ(roll angle) and α(pitch angle) as a function of time for bee # 5. The profiles shown in red, green, magenta, black and blue represents Θ_X_, Θ_Y_, Θ_Z_, θ and α.

**Figure 17:**
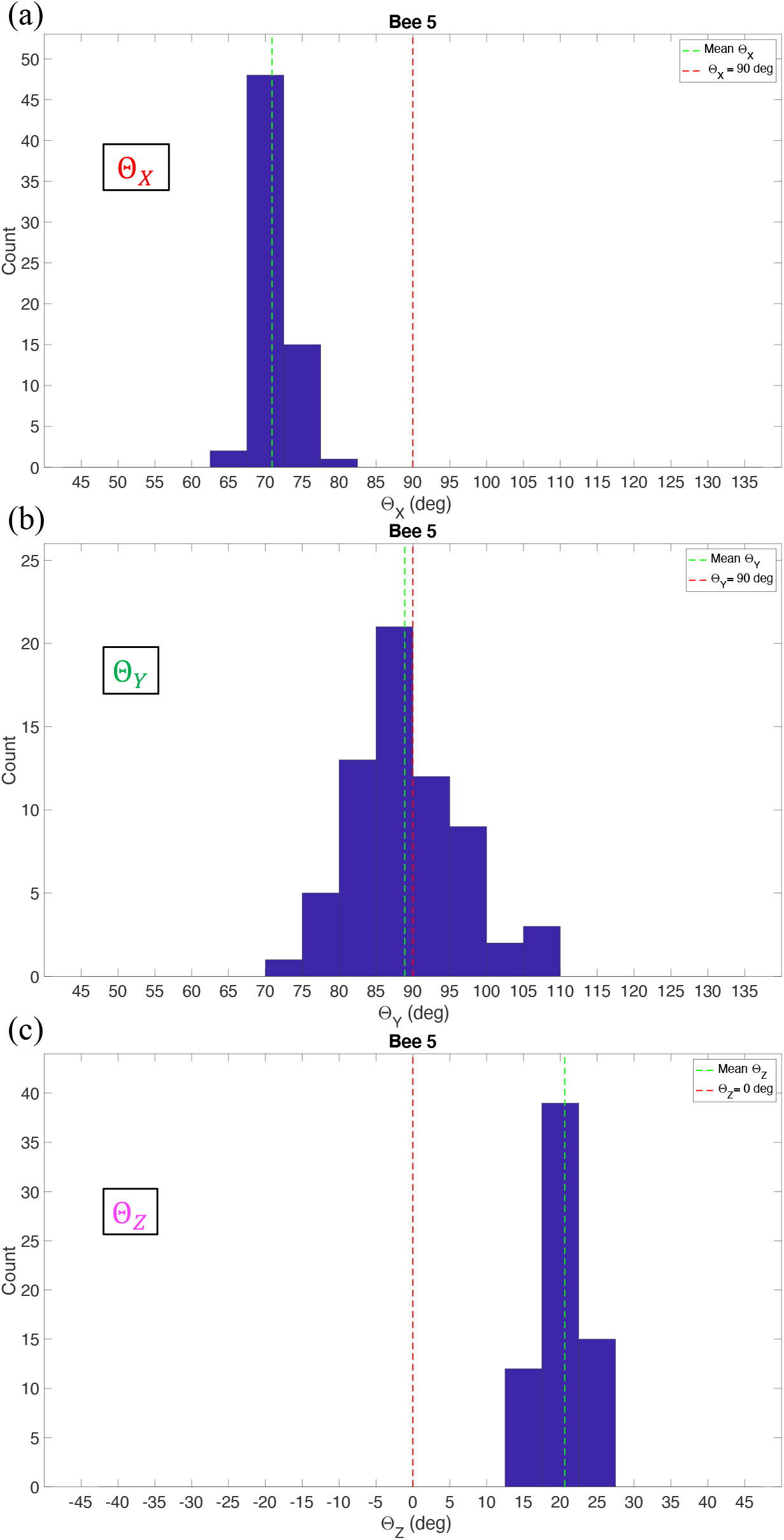
Histogram of Θ_X_, Θ_Y_, Θ_Z_ for the flight segment of bee #5.

Histograms of *Θ*_*X*_, *Θ*_*Y*_, and *Θ*_*Z*_, and the magnitudes and directions of the TFV for each of the 37 tunnel flight segments are given in Supplementary material (see Figs. S2). Table 3 shows the grand means of the various parameters that were measured and computed for these 37 flight segments.

**Table 3:** Grand means and standard deviations of Θ_X_, Θ_Y_, Θ_Z_, pitch angle (α), roll angle, absolute roll angle, absolute curvature, flight speed, and the magnitude of the total force vector (TFV) for the 37 flight segments filmed in the tunnel.

| $\theta_x$<br>(deg) | $\theta_y$<br>(deg) | $\theta_z$<br>(deg) | $\alpha$<br>(deg) | Roll<br>angle<br>(deg) | Absolute<br>roll<br>angle<br>(deg) | Tangential<br>acceleration<br>(m/s<sup>2</sup>) | Curvature<br>magnitude<br>(m<sup>-1</sup>) | Flight<br>speed<br>(m/s) | TFV<br>magnitude<br>(N) |
| --- | --- | --- | --- | --- | --- | --- | --- | --- | --- |
| 69.96<br>$\pm 6.66$ | 88.74<br>$\pm 4.80$ | 23.34<br>$\pm 2.72$ | 16.21<br>$\pm 4.78$ | 8.28<br>$\pm 14.55$ | 15.75<br>$\pm 9.45$ | $0.005 \pm 1.01$ | 13.60<br>$\pm 5.03$ | 0.55<br>$\pm 0.08$ | $(9.93 \pm 0.31) \times 10^{-4}$ |

The data in Table 3 confirm that (a) the magnitude of the TFV is maintained at a nearly constant value of (9.93±0.31) ×10^−4^ N; and (b) the TFV vector is maintained at a more or less constant orientation relative to the body, irrespective of substantial changes in the speed, curvature and attitude of flight. Furthermore, the TFV is always oriented almost exactly perpendicular to the bee’s lateral axis; it is not tilted to the left or the right, irrespective of the attitude or speed of flight, or the curvature of the flight trajectory.

## DISCUSSION

In this study we only analysed flights that were in or close to the horizontal plane. The reasons for this were (i) the cameras were pointed horizontally, and accurate measurement of roll angles was possible only for flights that were in the horizontal plane and close to the centre of the camera’s field of view; and (ii) the data on the wind-induced drag and lift generated by the body were measured only for horizontal flight conditions in the study by Nachtigall and Hanauer-Thieser^15^.

The essential findings of my study are summarized in Fig. 18. In the cloud environment, the mean magnitude of the TFV is 11.1×10^−4^ N. The TFV vector is oriented perpendicular to the body’s lateral axis, and at an elevation of 76.8° with respect to the body’s frontal axis. In the tunnel environment, the mean magnitude of the TFV is 9.9 ×10^−4^ N. The TFV vector is again oriented perpendicular to the body’s lateral axis, and at an elevation of 70° with respect to the body’s frontal axis. These numbers are more or less constant - regardless of the instantaneous speed, roll angle, or curvature of the flight trajectory.

**Figure 18:**
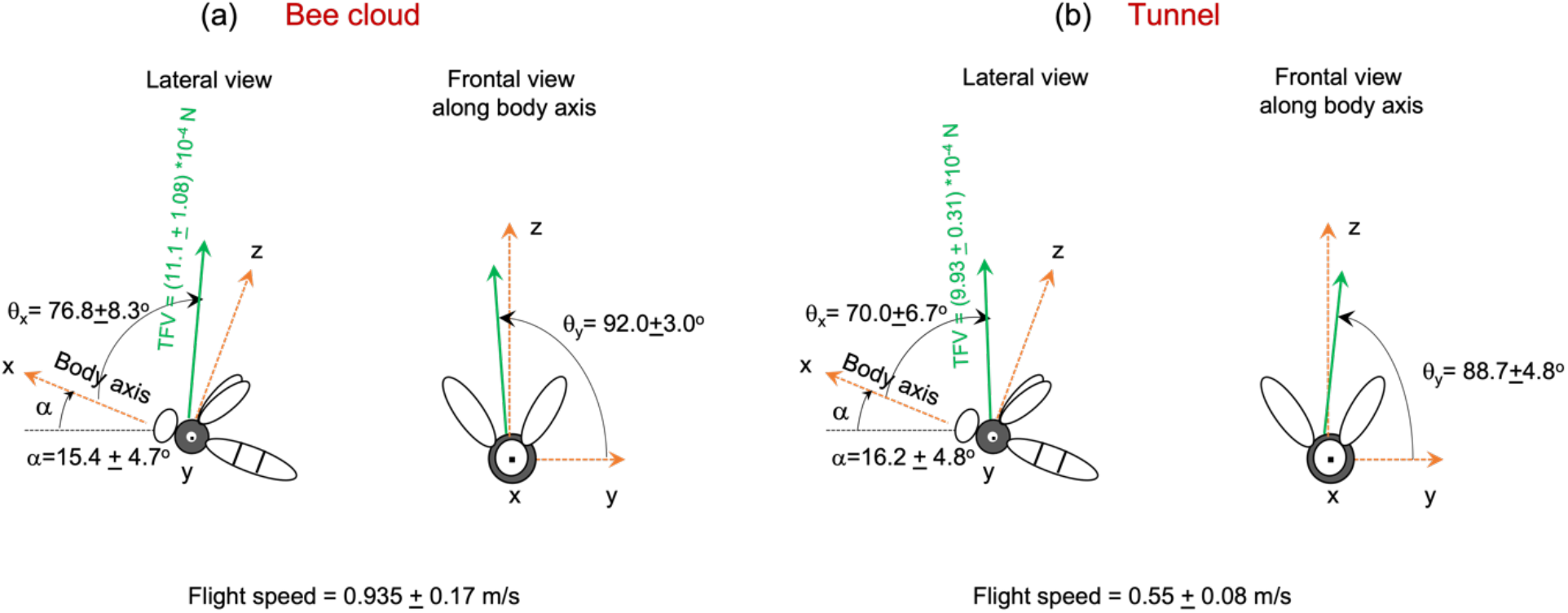
Estimated mean orientation of the total force vector (TFV) with respect to the bee’s body coordinates (x, y, z) for flights in the (a) bee cloud and (b) the tunnel. α is the pitch angle. The estimates (grand means and standard deviations) were obtained from analysing 27 flight segments in the bee could and 37 flight segments in the tunnel.

Even though the magnitudes and directions of the TFV estimated for the bees in the tunnel and the cloud are significantly different (TFV: *p =* 5.4269 × 10^−6^; *Θ*_*X*_: *p* = 0.0033; *Θ*_*Y*_: *p =* 1.1696 × 10^−4^; *Θ*_*Z*_: *p =* 3.8318 × 10^−4^; *two sample t-test*), the mean values differ only by small amounts. The pitch angles are not significantly different in the two environments (*p* = 0.71). However, the mean flight speed is significantly higher in the cloud environment (0.93 m/s) than in the tunnel environment (0.55 m/s), (*p* = 3.3760 × 10^−8^, *two sample t-test*).

Earlier studies have estimated the magnitude of the total flight force generated by *Drosophila melanogaster*^12,13^, *Musca domestica*^*12*^ *and Calliphora erythrocepha*^*18,19*^ under tethered conditions still air. The TFV magnitude was reported to be 3×10^−6^ N for *Drosophila*^*12*^ and 1.2 ×10^−4^ N for *Musca*^*12*^. Tethered *Calliphora erythrocephala* is reported to generate a TFV magnitude of about 9.5 ×10^−4^ N in still air [as we estimate from Fig. 4 of Blondeau^19^]. Tethered *Calliphora vicina* produces a lift of 3 ×10^−4^ N in an airstream of 1.5m/s [as we estimate from Fig. 4f of Nachtigall and Roth^18^]. Assuming a weight of 30mg for *Calliphora*^20^, which corresponds to a downward force of 0.29 ×10^−4^ N, it is evident that the lift force generated by *Calliphora* is comparable to, or slightly larger than its body weight.

My estimates of the mean magnitude of the TFV for the honeybee are 11.1 ×10^−4^ N (from the 27 flight segments in the cloud), and 9.9 ×10^−4^N (from the 37 flight segments in the tunnel), yielding an overall (weighted) mean magnitude of 10.4 ×10^−4^ N for the 64 flight segments that were analysed in this study. Since the honeybee is larger and heavier than *Drosophila* and *Musca*, it is reasonable to expect the TFV magnitude for the honeybee to be greater than those for flies.

The inclination of the TFV with respect to the frontal body axis was reported to be 24° for *Drosophila*^*12*^ and 29° for *Musca*^*12*^. The TFV inclinations that we have estimated for the honeybee are considerably higher: 76.8° in the cloud environment, and 70° in the tunnel environment. The reason for this discrepancy is not clear. One possibility is that tethering induces experimental artefacts. Indeed, this has been suggested by Fry et al^21^, who compared the forces generated by *Drosophila* in tethered and free-flight conditions. They found differences in the force profiles of the wingbeat cycles under the two conditions. They also noted that free-flying *Drosophila* hovers at a pitch angle of 45°. Since the TFV supports only the weight of the insect during hover (and not any accelerations or drag forces), their finding implies that, in freely-flying *Drosophila*, the TFV is inclined at an angle of (90° – 45°) = 45° with respect to the frontal body axis. This figure is considerably larger than the value of 24° reported in the earlier tethered-flight studies on *Drosophila*. Wagner^22^ measured the body pitch angles of freely-flying *Musca* at various flight speeds. He found that the body pitch increases as the flight speed decreases, reaching a value of about 40° at near-zero flight speeds. This implies that, in freely-flying *Musca*, the TFV is inclined at an angle of approximately (90° – 40°) = 50° with respect to the frontal body axis.

My finding that the total force vector maintains a more or less constant magnitude and orientation relative to the body axes - irrespective of various aerial manoeuvres in free flight - suggests that control of honeybee flight is described reasonably well by the so-called ‘helicopter’ model^23-25^. In a helicopter model, the tangential acceleration or deceleration along the flight trajectory would be controlled by lowering the pitch angle (to tilt the TFV forwards) or increasing the pitch angle (to tilt the TFV backwards). Thus, one would expect the pitch angle to be correlated with the tangential acceleration (or deceleration). Fig. 19 shows a scatterplot of tangential acceleration versus pitch angle, pooled across the 27 flight segments that were analysed in the cloud. Despite the considerable variability in the plot, there is a clear trend: lower pitch angles tend to be associated with acceleration, and higher pitch angles with deceleration. The linear regression displays a low correlation coefficient (r^2^ = 0.078), but a Fisher’s test reveals a highly significant trend (F = 311, *p* = 6.79 ×10^−67^). The regression line intersects the pitch axis at a value of 10.6 deg. This implies that, at this pitch angle, the bee would hover, or fly at a constant speed (neglecting body-induced drag), and that the TFV would be oriented vertically, because it is then only supporting the weight of the bee. Adding this pitch angle of 10.6 deg to the value of *Θ*_*X*_ (76.8 deg, see Fig. 18a) yields a result of 87.4 deg. This value is very close 90 deg, indicating that the TFV is indeed oriented approximately vertically in this condition. This adds further support to the accuracy of the estimate of *Θ*_*X*_. Part of the reason for the large variability in the tangential accelerations in the scatterplot may be the presence of turning phases in the trajectories, where the TFV (which is constant in magnitude) supports centripetal as well as tangential accelerations.

**Figure 19:**
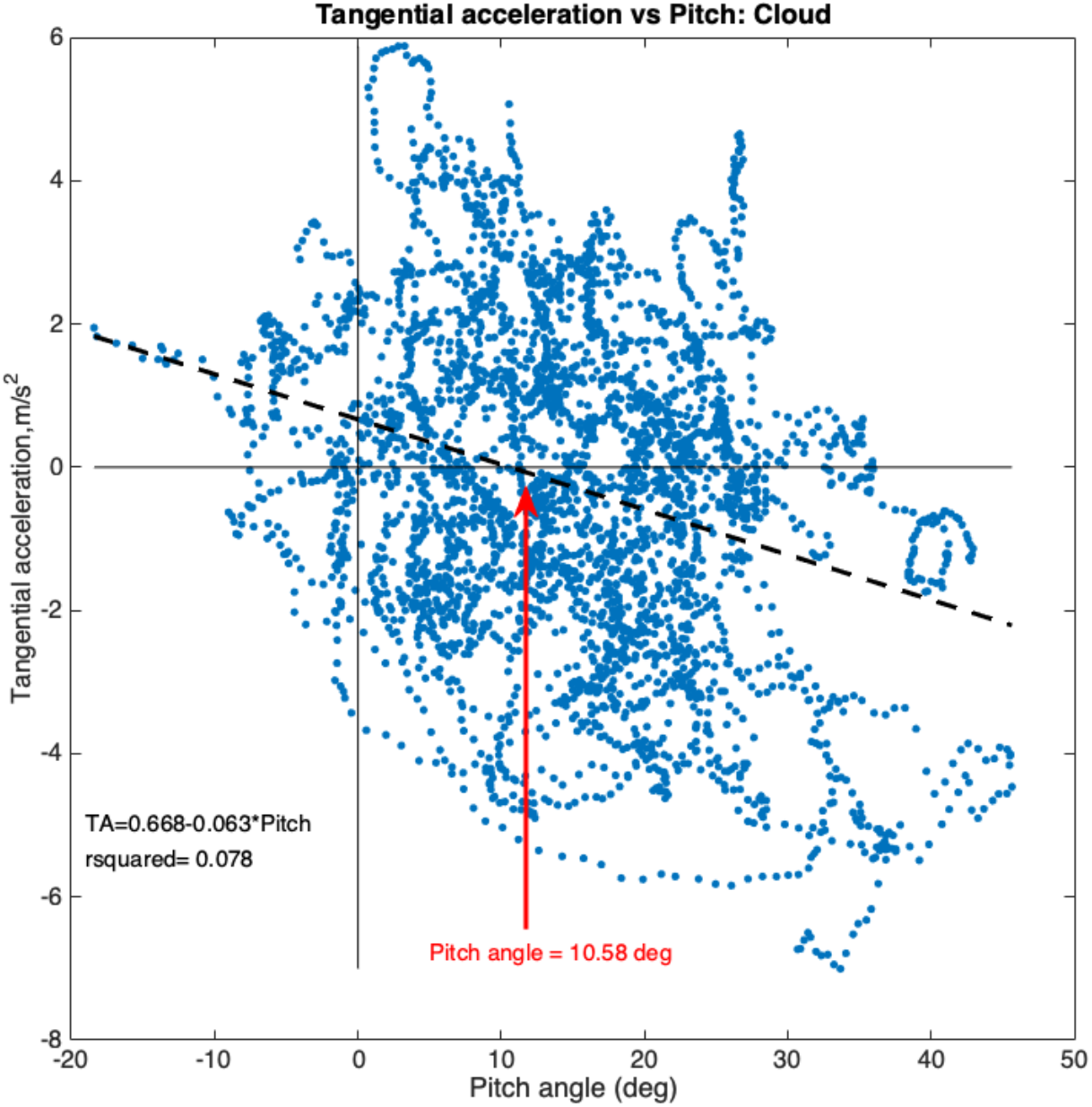
Scatterplot and linear regression of tangential acceleration versus body pitch angle pooled across the 27 flight segments that were analysed in the cloud. Details in text.

Fig. 20 shows a similar scatterplot of tangential acceleration versus pitch angle, pooled across the 37 flight segments that were analysed in the tunnel. The linear regression again displays a low correlation coefficient (r^2^ = 0.087), but a Fisher’s test reveals a highly significant trend (F = 683, *p* = 5.96×10^−144^). The regression line intersects the pitch axis at a value of 18.3 deg. The result of adding this pitch angle to the value of *Θ*_*X*_ (70.0 deg, see Fig. 18b) yields a result of 88.3 deg, which is again very close 90 deg, indicating that the TFV is again oriented approximately vertically in this condition, and supporting the validity of the estimate of *Θ*_*X*_for the tunnel flights.

**Figure 20:**
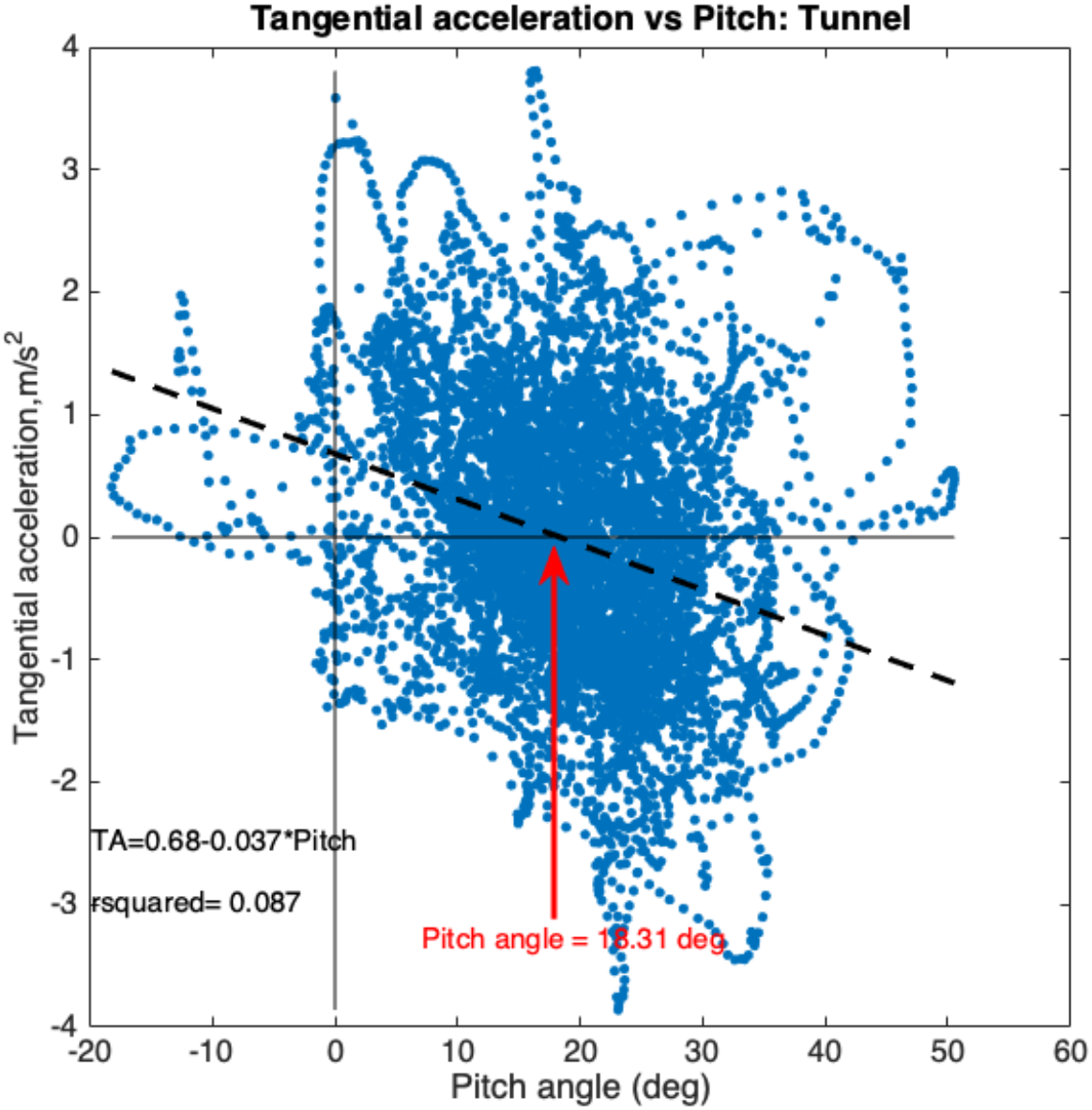
Scatterplot and linear regression of tangential acceleration versus body pitch angle pooled across the 37 flight segments that were analysed in the tunnel. Details in text.

The mean TFV magnitude of 10.4 ×10^−4^N for the honeybee is only slightly greater than the downward force (9.81×10^−4^N) due to its body weight (0.1g). This parallels the matching of the lift force and the body weight in the case of the blowfly *Calliphora vicina* (see above).

Collectively, the above findings reveal that honeybees generate a total flight force that is largely invariant in its magnitude, as well as its orientation relative to the body – in accordance with the ‘helicopter’ model. Accelerations and decelerations are achieved by decreasing the pitch of the body to tilt the TFV forwards to create a forward thrust component, or increasing the pitch of the body to tilt the TFV backwards to create a reverse thrust component. During a turn the centrifugal force experienced by the body is countered by rolling into the turn, which tilts the TFV into the turn to generate a centripetal force component. Of course, turns typically include a yaw motion, in addition to the roll. It is likely that yaw and roll are generated conjointly by increasing the amplitude of the wingbeat on the outside of the turn, and decreasing it on the inside^13,19,26^. This simultaneously increases the lift and thrust generated by the outside wing, (or outside pair of wings, in the case of the honeybee) and decreases the lift and thrust generated by the inside wing - thereby generating a roll torque as well as a yaw torque. Yaw motions are also aided by ruddering actions of the abdomen and the hind legs^27^. My findings do not, however, exclude the possibility of small changes in the direction of the TFV relative to the body - which could be implemented by varying the stroke plane of the wings, or other detailed features of wing articulation, as has been suggested by Fry et al^21^, and described in detail in the Introduction.

## CONCLUSIONS

The total flight force exerted by a bee is surprisingly constant, irrespective of the speed of flight or the attitude of the body. This force also has a constant orientation with respect to the bee’s body axes. These findings are true for flights in the curved tunnel, as well as those in the cloud. The results indicate that turns are executed (a) by holding the magnitude of the TFV constant and (b) by redirecting the TFV appropriately, through reorientation of the body, to achieve various flight manoeuvres.

